# Pleiotropic effects of BAFF on the senescence-associated secretome and growth arrest

**DOI:** 10.1101/2022.10.25.513730

**Authors:** Martina Rossi, Carlos Anerillas, Maria Laura Idda, Rachel Munk, Chang Hoon Shin, Stefano Donegà, Dimitrios Tsitsipatis, Allison B. Herman, Jennifer L. Martindale, Xiaoling Yang, Yulan Piao, Krystyna Mazan-Mamczarz, Jinshui Fan, Luigi Ferrucci, Supriyo De, Kotb Abdelmohsen, Myriam Gorospe

## Abstract

Senescent cells release a variety of cytokines, proteases, and growth factors collectively known as the senescence-associated secretory phenotype (SASP). Sustained SASP contributes to a pattern of chronic inflammation associated with aging and implicated in many age-related diseases. Here, we investigated the expression and function of the immunomodulatory cytokine BAFF (B-cell activating factor), a SASP protein, in multiple senescence models. We first characterized BAFF production across different senescence models, including senescent human diploid fibroblasts (WI-38, IMR-90) and monocytic leukemia cells (THP-1), and tissues of mice induced to undergo senescence. We then identified IRF1 (interferon response factor 1) as a transcription factor required for promoting *BAFF* mRNA transcription in senescence. We discovered that suppressing BAFF production decreased the senescent phenotype of both fibroblasts and monocyte-derived THP-1 cells, overall reducing IL6 secretion, SA-β-Gal staining, and γ-H2AX accumulation. Importantly, however, the influence of BAFF on the senescence program was cell type-specific: in monocytes, BAFF promoted the early activation of NF-κB and general SASP secretion, while in fibroblasts, BAFF contributed to the production and function of TP53 (p53). We propose that BAFF is elevated across senescence models and is a potential target for senotherapy.

## Introduction

Cellular senescence is a state of indefinite cell cycle arrest arising in response to a variety of stressful stimuli, including telomere erosion, oncogenic signaling, and damage to DNA and other molecules (Hayflick 1965; Munoz-Espin and Serrano 2014). Despite experiencing persistent growth arrest, senescent cells remain metabolically active and express and secrete distinct subsets of proteins, including cytokines, chemokines, metalloproteases, and growth factors, a trait collectively known as the senescence-associated secretory phenotype (SASP). The sustained production of SASP factors in tissues and organs promotes the recruitment of immune cells, tissue remodeling, and chronic inflammation at the systemic level (Basisty et al. 2020; Franceschi and Campisi 2014). Senescent cells are necessary for developmental processes like embryonic morphogenesis and wound healing, and have tumor-suppressive properties in young individuals. However, with advancing age, the accumulation of senescent cells exacerbates age-related pathologies like cancer, diabetes, and neurodegenerative and cardiovascular diseases (McHugh and Gil 2018; Munoz-Espin and Serrano 2014; Franceschi et al. 2007).

Despite the recognized impact of senescent cells, the unequivocal detection of senescent cells remains a challenge. Several markers of senescence have been described in cultured cells as well as in tissues and organs, but they are not universal markers of all senescent cells, and they are often not exclusive of senescent cells, as non-senescent cells may express them as well. Therefore, multiple markers must be present in order to identify a cell as senescent. Classic markers of senescence include cell cycle inhibitors [e.g., the cyclin-dependent kinase (CDK) inhibitor proteins p16 (CDKN2A) and p21(CDKN1A)], the presence of nuclear foci of unresolved DNA damage (often visualized by the presence of a phosphorylated histone, γ-H2AX), and increased activity of a senescence-associated β-Galactosidase (SA-β-Gal) that functions at ~pH 6 (Gorgoulis et al. 2019; Hernandez-Segura, Nehme, and Demaria 2018; Idda et al. 2020). Our recent screen identified the *TNFSF13B* (*Tumor Necrosis Factor Superfamily, Member 13B*) mRNA, encoding the cytokine BAFF (B cell-activating factor), as an RNA marker shared across a number of senescent cell types and inducers (Casella et al. 2019). BAFF is produced as a membrane-bound or secreted cytokine that plays an essential role in the homeostasis of the immune system (Kalled 2005; Mackay and Schneider 2009; Rauch et al. 2009; Smulski and Eibel 2018). However, BAFF also plays a key role in sustaining the inflammatory processes associated with autoimmune diseases like systemic lupus erythematosus, multiple sclerosis, and rheumatoid arthritis (Idda et al. 2019; Kalled 2005; Steri et al. 2017). The pro-inflammatory role of BAFF is primarily elicited by the activation of three receptors [BAFFR (TNFRSF13C), TACI (TNFRSF13B) and BCMA (TNFRSF17)], which converge on paths that signal through the transcription factor NF-κB (Nicoletti et al. 2016; Eslami and Schneider 2021; Bossen and Schneider 2006; Smulski and Eibel 2018). In turn, NF-κB transcriptionally activates the production of many pro-inflammatory and adhesion factors (Tang, Zhang, and Wei 2018). However, the role of BAFF in cell senescence is unknown.

Here, we investigated the expression and function of BAFF in senescence. We present evidence that BAFF is elevated in models of cell senescence in mice and cultured human cells. In response to a range of inducers, the levels of *BAFF* mRNA and total cellular BAFF were increased, as were the levels of secreted BAFF in the culture media of senescent cells and in the blood of mice following doxorubicin-induced senescence. Mechanistically, the transcription factor IRF1 (interferon response factor 1) was found to increase *BAFF* mRNA levels in senescent cells via activated transcription. In the presence of secreted BAFF, monocyte-derived cells activated NF-κB, which in turn transcriptionally induced the production of SASP factors, while senescent fibroblasts instead activated a p53-dependent gene expression program. Our data indicate that BAFF promotes senescence in a pleiotropic manner, enhancing the SASP in some cell types and p53-mediated growth arrest in others.

## Results

### BAFF increases in cultured senescent cells and in a senescent mouse model

Our previous analysis of multiple models of senescence, including human primary fibroblasts (WI-38, IMR-90), umbilical vein endothelial cells (HUVECs), and aortic endothelial cells (HAECs), revealed a unique senescence transcriptome signature (Casella et al. 2019), including heightened production of the *TNFSF13B* mRNA, encoding the cytokine BAFF, a SASP factor. RNA sequencing (RNA-seq) analysis (Casella et al. 2019) indicated that the levels of *TNFSF13B/BAFF* mRNA were elevated across all the senescence models tested previously. These models included WI-38 fibroblasts that were proliferating at population doubling level (PDL) 25 and then rendered replicatively senescent (RS) by division until exhaustion (at ~PDL55), exposed to ionizing radiation (IR at 10 Gy, evaluated 10 d later), infected for 18 h with a lentivirus that triggered oncogene-induced senescence (OIS) by expression of HRAS^G12V^ and evaluated 8 d later, or treated with doxorubicin (DOX) for 24 h and evaluated 7 d later. Additional models tested included proliferating IMR-90 fibroblasts (PDL25) rendered RS by culture to ~PDL55 or senescent by exposure to IR (10 Gy, assessed 10 d later), and HUVECs and HAECs which were either proliferating or rendered senescent by exposure to IR (4 Gy, evaluated 10 d later) (**Figure 1—figure supplement 1A**).

Given that BAFF is mainly expressed by myeloid cells like monocytes and dendritic cells (Sakai and Akkoyunlu 2017; Steri et al. 2017), we extended the analysis of BAFF expression in senescence using the human monocytic leukemia-derived cell line THP-1. First, we induced senescence in THP-1 cells by treatment with IR (5 Gy for 6 days, a dose selected because it suppressed cell growth but did not reduce cell viability, **Figure 1—figure supplement 1B,C**). Similarly, we selected 10 nM DOX treatment for 48 h, followed by 4 additional days in DOX-free media, as a treatment dose that induced senescence in THP-1 cells without reducing viability (not shown). As additional models of senescence, we included primary WI-38 fibroblasts rendered senescent by exposure to IR or to HRAS^G12V^ (OIS) as described above (**Figure 1A-D**). In these treatment groups, we first found markedly increased senescence-associated (SA)-β-Gal levels in all four senescence groups (Methods; **Figure 1A**). We then quantified the levels of *BAFF* mRNA by reverse transcription (RT) followed by real-time, quantitative (q)PCR analysis and normalized them to the levels of *ACTB* mRNA, encoding the housekeeping protein ACTB (β-Actin) (Methods; **Figure 1B**); as shown, *BAFF* mRNA levels increased markedly in all four senescence groups.

**Figure 1.**
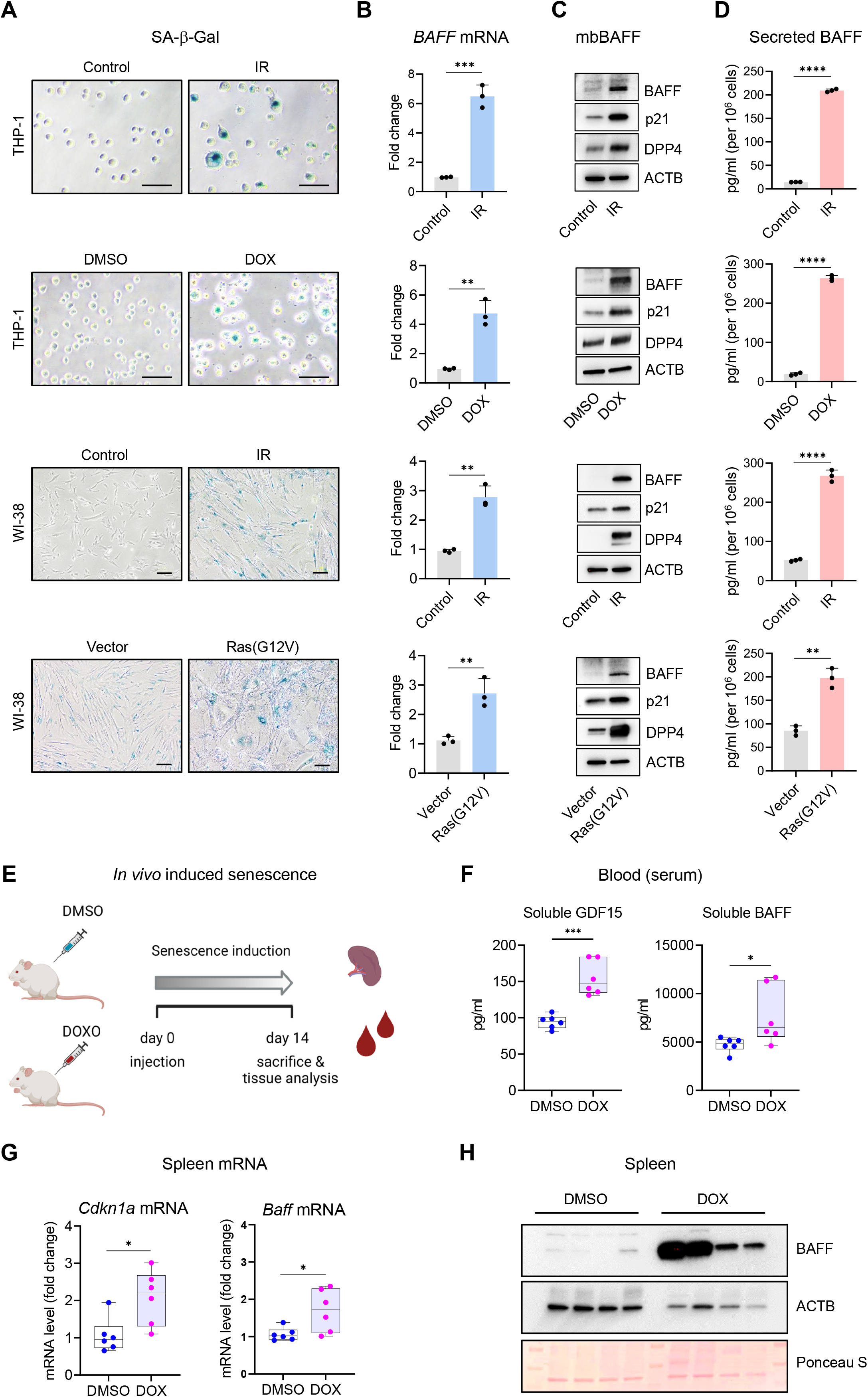
Increased BAFF expression in cultured senescent cells and in mouse model of senescence. (**A**) Micrographs of SA-β-Gal activity in senescent cultured cells relative to corresponding proliferating controls. From top: THP-1 cells untreated (Control, proliferating) or treated with a single dose of IR (5 Gy) and cultured for an additional 6 days; THP-1 cells treated with DMSO or with a single dose of 10 nM doxorubicin (DOX) for 48 h, followed by 4 additional days in culture without DOX (for a total of 6 days since the treatment with DOX); WI-38 fibroblasts that were proliferating or had been treated with a single dose of IR (10 Gy) IR and cultured for an additional 10 days; WI-38 fibroblasts transduced for 18 h with a control lentivirus (empty vector) or with a RAS(G12V) lentivirus, whereupon culture medium was replaced and cells were cultured for an additional 8 days. (**B**) RT-qPCR analysis of the levels of *BAFF* (*TNFSF13B*) mRNA, normalized to the levels of *ACTB* mRNA [encoding the housekeeping protein ACTB (β-Actin)] in cells processed as in (A). (**C**) Western blot analysis of the levels of membrane-bound BAFF (mbBAFF), DPP4, p21, and loading control ACTB in cells processed as in (A). (**D**) Levels of BAFF secreted in the culture media in cells treated as described in (A), as quantified by ELISA. (**E**) Schematic of the strategy to induce senescence in DOX-treated mice (Materials and methods), created with BioRender. (**F**) Quantification of the levels of soluble GDF15 and BAFF in the blood (serum) of mice treated as in (E) using ELISA (Materials and methods). GDF15 was used as a positive control of induced senescence (Basisty et al., 2020). (**G**) RT-qPCR analysis of the levels of *Cdkn1a* and *Baff* mRNAs in spleens of mice treated as in (E). *18S* rRNA levels were also quantified and served to control for loading. (**H**) Representative western blot analysis of the levels of BAFF in spleen homogenates obtained from mice treated as in (E). ACTB and Ponceau S staining served as loading controls. Significance (*p < 0.05, **p < 0.01, ***p < 0.001, ****p <0.0001) was assessed with Student’s *t*-test in all panels. Scale bars, 100 μm. Source Data for Figure 1: **Figure 1—Source Data 1.** Uncropped immunoblots for Figure 1.

We subsequently measured BAFF protein levels by western blot analysis alongside senescence protein markers DPP4 and p21, known to increase during senescence, and loading control ACTB (**Figure 1C**). Finally, given that BAFF exerts its function as a membrane-bound protein and a secreted cytokine (Eslami and Schneider 2021), we measured BAFF in conditioned media using ELISA; as shown (**Figure 1D**), the levels of secreted BAFF increased in conditioned media from senescent cells. The levels of *BAFF* mRNA and secreted BAFF were also measured in other models of senescence, as confirmed by seeing enhanced SA-β-Gal expression in IMR-90 fibroblasts treated with etoposide (ETO, 50 μM for 8 days), WI-38 fibroblasts rendered senescent by RS or ETO, and human vascular smooth muscle cells (hVSMCs) exposed to 10 Gy IR and cultured for an additional 7 days (**Figure 1—figure supplement 1D**). In all cases, *BAFF* mRNA levels increased during senescence (**Figure 1—figure supplement 1E**), and secreted BAFF was generally elevated with senescence, although it was undetectable in hVSMCs, and the levels were overall higher in THP-1 cells (**Figure 1D and Figure 1— figure supplement 1F**). In addition, we confirmed the increased expression of cellular BAFF by immunofluorescence (red, BAFF; blue, DAPI-stained nuclei) in senescent (IR) relative to proliferating cells in both THP-1 cells and WI-38 cells (**Figure 1—figure supplement 1G,H**).

Next, we sought to evaluate the levels of BAFF expressed in mice in which senescent cells were induced to accumulate in tissues and organs. We triggered a rise in senescent cells in mice by injecting DOX (10 mg/kg) once, and measuring BAFF levels in the serum 14 days later (**Figure 1E** and Materials and methods). We observed a modest but significant increase in the levels of circulating BAFF (**Figure 1F**), with a difference greater than 1600 pg/ml in the median between the two groups. As a positive control that senescence was induced in the mouse, we measured the increase in the circulating levels of the SASP core factor GDF15 (**Figure 1F**) (Basisty et al. 2020). Next, we analyzed the expression of BAFF in the spleen, a major reservoir of myeloid cells. We observed a significant increase in the levels of *Baff* mRNA, as measured by RT-qPCR analysis; we measured the levels of senescence marker *Cdkn1a* (*p21*) mRNA alongside as a positive control (**Figure 1G**). To study if the elevation in *Baff* mRNA levels yielded increased BAFF protein production, we performed western blot analysis on homogenates from spleen and observed high levels of BAFF protein in DOX-treated mice compared to control (DMSO-treated) mice (**Figure 1H**). Given the high variability in ACTB expression among different mice, we also used Ponceau S staining to monitor loading differences in western blots (**Figure 1H**). Together, these results indicate that BAFF is elevated in senescent cultured cells and also in senescent cells *in vivo*.

### IRF1 promotes *BAFF* mRNA transcription in senescence

Next, we investigated if the rise in *BAFF* mRNA with senescence in THP-1 cells was the result of transcriptional or posttranscriptional regulatory mechanisms. In THP-1 cells rendered senescent by exposure to IR, we assessed the changes in levels of *BAFF* pre-mRNA (a surrogate for *de novo* transcription) by RT-qPCR analysis and found that they mirrored those of total *BAFF* mRNA (**Figure 2A**). These results indicated that the rise in *BAFF* mRNA was strongly driven by increased transcription. The notion that the rise in *BAFF* mRNA levels was the result of increased transcription and not increased *BAFF* mRNA stability was directly tested by measuring the half-lives of *BAFF* mRNA after adding the transcriptional inhibitor Actinomycin D; as shown in **Figure 2—figure supplement 1A**, the rate of *BAFF* mRNA decay in proliferating THP-1 cells (t_1/2_~4 h) was comparable to that observed in THP-1 cells rendered senescent by IR (t_1/2_~3 h), indicating that *BAFF* mRNA was not longer-lived in senescent THP-1 cells, and further supporting the idea that *BAFF* mRNA increased through elevated transcription. Similarly, WI-38 cells rendered senescent by RS or IR showed increased *BAFF* pre-mRNA levels that mirrored the rise in *BAFF* mRNA and *BAFF* mRNA had comparable half-lives across the proliferating and senescent populations (**Figure 2—figure supplement 1B-D**), suggesting that *BAFF* mRNA was transcriptionally elevated across senescence models. In mRNA stability assays, the unstable *MYC* mRNA was measured as an internal control (**Figure 2—figure supplement 1A,C,D**).

**Figure 2.**
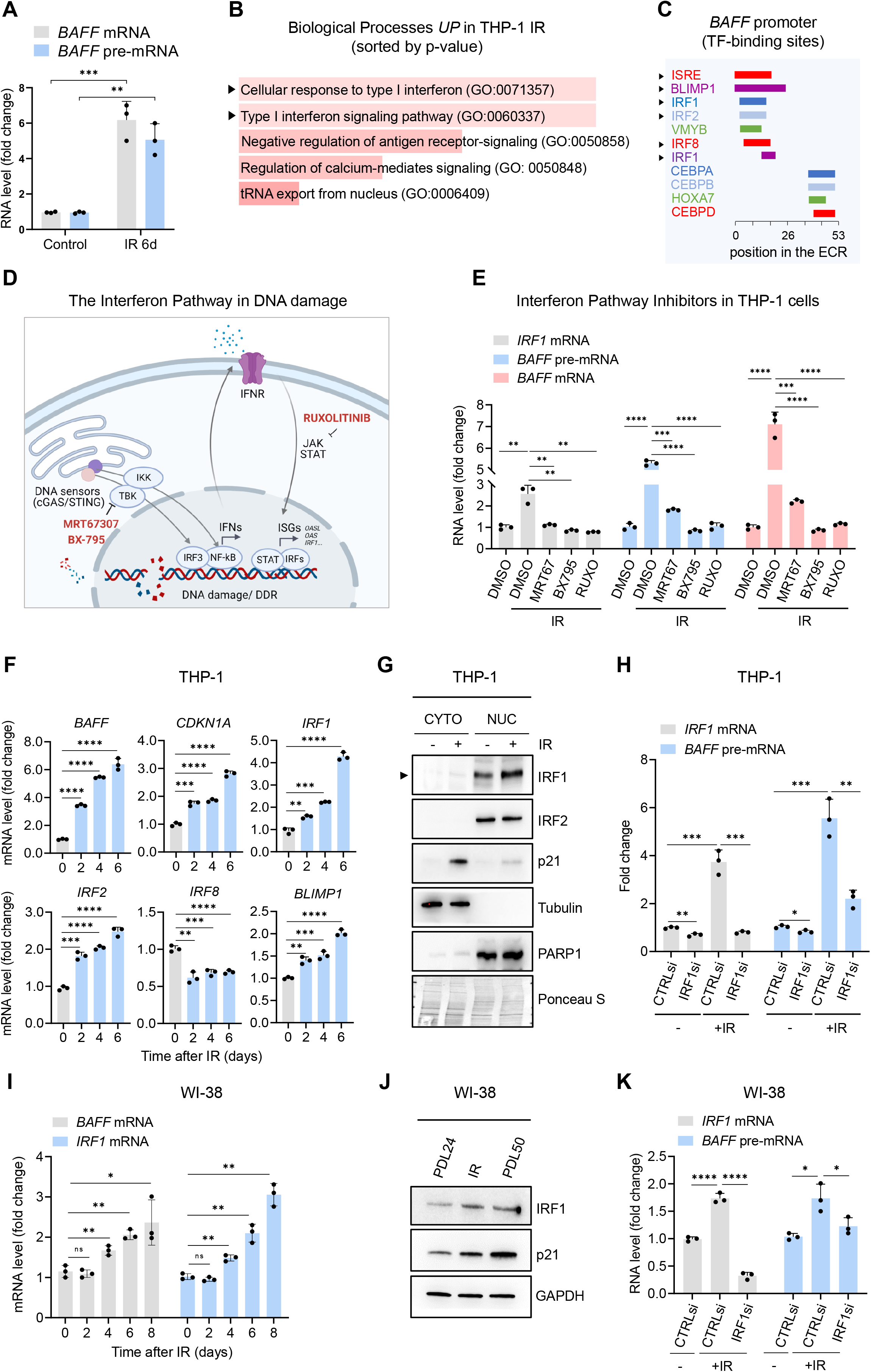
Interferon response pathway and IRF1 promote *BAFF* transcription in senescence. (**A**) RT-qPCR analysis of the levels of *BAFF* mRNA and *BAFF* pre-mRNA in THP1 cells that were either untreated (Control) or treated with IR (5 Gy) and analyzed 6 days later. (**B**) Terms enriched by Gene Ontology analysis after proteomic analysis of THP-1 cells that were rendered senescent by IR relative to untreated control cells. Proteomic analysis is available as **Figure 2-Source Data 1**. GO terms were sorted by p-value ranking (EnrichR analysis). (**C**) Schematic of conserved TF binding sites on the *BAFF* promoter, as analyzed using ECR browser and rVista 2.0. (**D**) Schematic of the interferon response triggered by DNA damage, highlighting the targets affected by the different interferon inhibitors used in our study (Hartlova et al. 2015; Li and Chen 2018; Fu et al. 2020); created using BioRender. (**E**) RT-qPCR analysis of the levels of *BAFF* mRNA, *BAFF* pre-mRNA, and *IRF1* mRNA in THP-1 cells after treatment with the inhibitors of the interferon pathway indicated in (D) [Ruxolitinib (Ruxo, 1 μM), MRT67307 (MRT67, 5 μM), and BX-795 (5 μM)] or with control vehicle DMSO for 5 days after treatment with IR. *IRF1* mRNA was included as a positive control of interferon-stimulated mRNAs. (**F**) RT-qPCR analysis of the levels of expression of *BAFF* and *p21* mRNAs and other interferon-regulated transcripts (*IRF1, IRF2, IRF8*, and *BLIMP1* mRNAs) in THP-1 cells that were left untreated or were treated with IR and cultured for the indicated times. (**G**) Western blot analysis of the levels of IRF1 and IRF2 in the cytoplasmic and nuclear fractions of THP-1 cells that were left untreated or were irradiated and assayed 6 days later. The cytoplasmic protein tubulin, the nuclear protein PARP1, and the senescence-associated protein p21 were included in the analysis. Ponceau staining of the transfer membrane was included to monitor differences in loading and transfer among samples. (**H**) RT-qPCR analysis of the levels of *IRF1* mRNA and *BAFF* pre-mRNA in THP-1 cells transfected with control (CTRLsi) or IRF1-directed (IRF1si) siRNAs 72 h after either no treatment (-) or treatment with IR. **(I)**RT-qPCR analysis showing the levels of *BAFF* and *IRF1* mRNAs in WI-38 cells treated with IR (10 Gy) and collected at the indicated times. (**J**) Western blot analysis of the levels of BAFF protein in WI-38 fibroblasts that were proliferating (PDL24), senescent following exposure to IR (10 Gy) and cultured for 10 days, and senescent through replicative exhaustion (PDL50); p21 was used as positive marker for senescence and GAPDH as loading control. (**K**) RT-qPCR analysis of the levels of *BAFF* pre-mRNA in WI-38 cells transfected with CTRLsi or IRF1si and either left untreated (-) or treated with IR (10 Gy) and assayed 5 days later. Significance (ns, not significant; *p < 0.05, **p < 0.01, ***p < 0.001, ****p <0.0001) was assessed with Student’s *t*-test. Source Data Files for Figure 2: **Figure 2—Source Data 1.** Proteomic analysis performed in control THP-1 cells treated with IR. **Figure 2—Source Data 2.** Uncropped western blot images for Figure 2.

To begin to identify the transcription factors (TFs) that might upregulate *BAFF* mRNA transcription in senescence, we analyzed subsets of proteins identified as changing, based on proteomic analysis, in THP-1 cells primed for senescence after irradiation with 5 Gy and culture for 72 h (**Figure 2—Source Data 1**). The proteomic analysis (**Figure 2B and Figure 2—Source Data 1**) showed a robust and predominant activation of the Type I interferon response pathway in IR-treated THP-1 cells. Importantly, DNA damage was previously shown to prime both the interferon response and inflammation (Hartlova et al. 2015; Li and Chen 2018). Computational analysis of the evolutionarily conserved regions (ECRs) in the *BAFF* promoter and the rVista2.0 database (https://ecrbrowser.dcode.org/) indicated the presence of an interferon-sensitive response element (ISRE) and multiple binding sites for interferon regulatory factors (IRFs), including IRF1, IRF2, and IRF8 (**Figure 2C**). To test the possible role of these TFs in driving *BAFF* transcription, we evaluated three inhibitors [Ruxolitinib (Ruxo), MRT67307 (MRT67), and BX-795] that target the interferon response pathway at different levels (**Figure 2D**), and measured the efficiency of these inhibitors by quantifying the levels of *IRF1* mRNA, a transcript that is inducible during the IFN response (**Figure 2D**) (Fujita et al. 1989; Forero et al. 2019; Panda et al. 2019). As shown, these inhibitors decreased the levels of *BAFF* pre-mRNA and *BAFF* mRNA in IR-induced senescent THP-1 cells; they also decreased *IRF1* mRNA levels, included here as a positive control (**Figure 2E**). These results uncovered a key role for the interferon response in promoting BAFF expression following senescence-inducing DNA damage and strengthened the hypothesis that IRFs enhanced BAFF transcription.

To narrow down our list of IRF candidates possibly involved in BAFF transcription (**Figure 2C**), we focused on those IRF factors whose expression by mRNA and protein either increased or remained unchanged (but did not decline) after senescence-inducing DNA damage. RT-qPCR analysis at the times shown following treatment of THP-1 cells with IR (5 Gy) revealed increased levels of *BAFF* and *CDKN1A* mRNAs, as well as increased levels of *IRF1, IRF2, BLIMP1* mRNAs and reduced *IRF8* mRNA levels (**Figure 2F**). Given that *IRF8* expression was strongly reduced in senescence and BLIMP1 is a known transcriptional repressor (Keller and Maniatis, 1991; Martins and Calame, 2008) we focused on IRF1 and IRF2 as potential inducers of *BAFF* mRNA transcription in senescent cells. THP-1 cell fractionation followed by analysis of changes in their subcellular distribution during senescence revealed elevated nuclear localization of IRF1, but not IRF2, in senescent cells (**Figure 2G**), suggesting that IRF1 might be primarily implicated in the transcription of *BAFF* mRNA. To test this possibility, in THP-1 cells rendered senescent by IR we silenced IRF1 using IRF1-specific siRNAs. RT-qPCR analysis confirmed the silencing of IRF1 and showed a significant reduction in *BAFF* pre-mRNA levels (**Figure 2H**). Similarly, the levels of *IRF1* mRNA and IRF1 protein increased in senescent WI-38 cells (**Figure 2I,J**), and silencing IRF1 decreased *BAFF* pre-mRNA levels (**Figure 2K**). Together, these findings support a role for the interferon pathway, and TF IRF1 in particular, in increasing the transcription of *BAFF* mRNA in senescent cells.

### Transcriptomic and proteomic analyses reveal BAFF-dependent roles in senescence

To identify the biological processes regulated by BAFF during senescence in monocytes, we performed transcriptomic and proteomic analyses in senescent THP-1 cells that expressed either normal or reduced levels of BAFF. Given that proliferating THP-1 cells express detectable levels of BAFF, and that BAFF rapidly increases in the first few days after IR treatment (**Figures 1C and 2F**), we silenced BAFF using siRNAs directed at *BAFF* mRNA in proliferating THP-1 cells, and then induced IR treatment the next day (**Figure 3A**). RNA-seq analysis (GSE213993, reviewer token mbubswukttkhtsf) performed 72 h after IR revealed 500 transcripts that were highly upregulated by IR but repressed in BAFF-silenced cells, suggesting that induction of these transcripts was dependent on the presence of BAFF (**Figure 3B and Figure 3—Source Data 1**). Molecular Signatures Database (MSigDB) analysis of these 500 transcripts revealed enrichments in pro-inflammatory gene sets, including those implicated in TNF signaling, NF-κB activation, the IL6/JAK/STAT3 pathway, and the interferon response (**Figure 3C**). Gene ontology (GO) ‘Biological process’ analysis (**Figure 3D**) revealed enrichments in pro-inflammatory and immune biological processes, including the inflammatory response and leukocyte activation. The top-ranked categories of GO ‘Molecular function’ included NAD+ nucleosidase activity and pattern receptor activity transcripts encoding Toll-like receptor components (TLR1, TLR2, TLR4) that play key roles in the innate immune system and inflammation (Mukherjee, Karmakar, and Babu 2016). GO ‘Cellular component’ analysis showed an enrichment in secretory granules and secretory vesicles, suggesting that BAFF could be involved in inflammatory and secretory pathways in senescence. Complementing this notion, transcripts encoding cytokines [IL1B (IL-1β), CCL2, TNFAIP6/TSG-6], chemokines (CXCL1, CXCL2, CXCL8), and alarmins (S100A8, S100A9) were less abundant in BAFF-silenced cells, as assessed in the RNA-seq dataset (**Figure 3E**), and validated by RT-qPCR analysis (**Figure 3F**). Interestingly, analysis of mRNAs that were reduced in irradiated cells and elevated in BAFF-depleted cells further suggested a role for BAFF in membrane transport (**Figure 3— figure supplement 1A,B and Figure 3—Source Data 1**).

**Figure 3.**
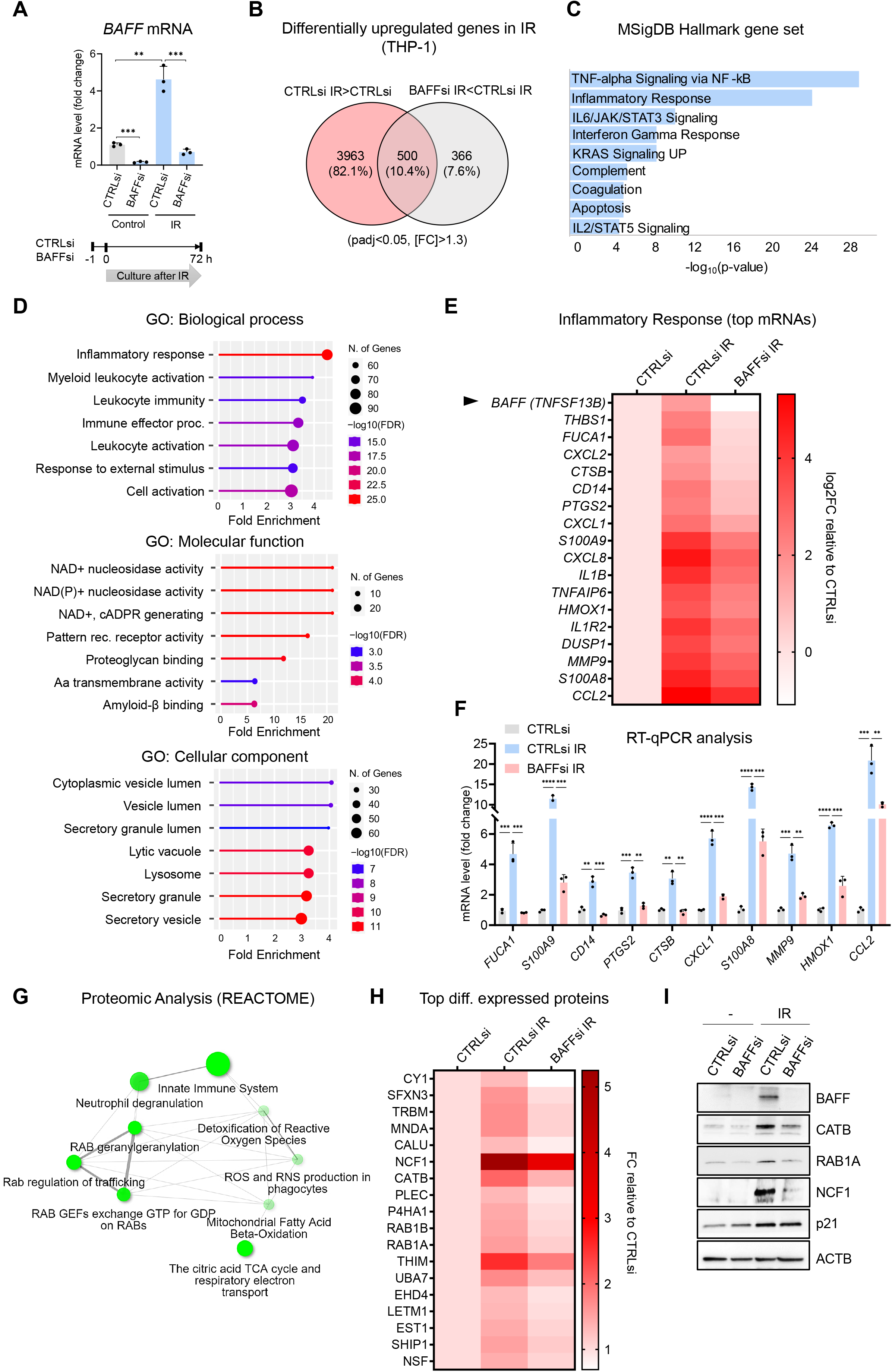
Transcriptomic and proteomic analysis in THP-1 cells suggests a role for BAFF in senescence-associated inflammation. (**A**) RT-qPCR analysis of the levels of *BAFF* mRNA in THP-1 cells transfected with CTRLsi or BAFFsi, irradiated 18 hr later (5 Gy), and assessed 72 h after that. *Bottom*, schematic of the timeline of BAFF silencing and exposure to IR in THP-1 cells. (**B**) Venn diagram of mRNAs identified by RNA-seq analysis as being differentially upregulated in THP-1 cells transfected with a control siRNA (CTRLsi) or BAFF siRNA (BAFFsi) following exposure to IR (5 Gy, collected 72 h later). Red circle: mRNAs upregulated in CTRLsi exposed to IR (CTRLsi IR) relative to non-irradiated cells (CTRLsi). Gray circle: mRNAs less induced in BAFFsi cells exposed to IR (BAFFsi IR) relative to CTRLsi exposed to IR (CTRLsi IR). A complete list of genes from the RNA-seq analysis is available (GSE213993, reviewer token mbubswukttkhtsf, and **Figure 3-Source Data 1**); cutoffs in Source Data: padj<0.05, [fold change] >1.3. (**C**) Molecular Signatures Database (MSigDB) hallmark analysis performed on the differentially expressed genes obtained from (B); diagram was created by EnrichR analysis and gene sets ordered by *p* value. (**D**) Gene ontology analysis (Biological Processes, Molecular Function and Cellular Component) performed on the differentially upregulated mRNAs identified in (B). (**E**) Heatmap of the differentially upregulated genes identified in (B) and included in the GO terms ‘Inflammatory Response’, ‘Leukocyte activation’, and ‘Immune effector process’, as well as those present in the MSigDB Hallmark term ‘Inflammatory Response’. Values are averages of duplicates. Top genes upregulated in IR were sorted according to their greater reduction after BAFF silencing. Data are shown as Log2FC in gene expression relative to untreated cells (CTRLsi: log2FC=0). (**F**) RT-qPCR analysis of a subset of differentially expressed mRNAs identified in (B) and (E). Significance (*p < 0.05, **p < 0.01, ***p < 0.001, ****p <0.0001) was assessed with Student’s *t*-test. **(G)** Reactome network analysis showing the most highly enriched categories of proteins differentially upregulated in THP-1 cells transfected with control (CTRLsi) or BAFF-directed (BAFFsi) siRNAs and the next day exposed to 5 Gy IR or left untreated and collected 72 h later (complete proteomic datasets are in **Figure 3-Source Data 2;** Cutoff fold change was 1.3). Two pathways (nodes) are connected if they share 10% or more proteins. Darker nodes are more significantly enriched protein sets; larger nodes represent larger protein sets. Thicker edges represent more overlapped proteins. **(H)** Heatmap of the top differentially upregulated proteins between CTRLsi and BAFFsi. Top proteins increased after IR were sorted according to their greater reduction after BAFF silencing. Data are shown as fold change between PSM (peptide spectrum matches) relative to the untreated (CTRLsi: FC=1). Cutoff: [FC] >1.3, protein with PSM above 15). A complete list of differentially upregulated proteins is available in **Figure 3-Source Data 2**. (**I**) Western blot analysis of representative proteins identified from the proteomic analysis in (G). p21 was included as a control for senescence and ACTB as a loading control. Diagrams in (D,G) were created with ShinyGO. Source Data Files for Figure 3: **Figure 3—Source Data 1.** RNA-seq analysis performed in THP-1 cells transfected with CTRLsi or BAFFsi and treated with IR. **Figure 3—Source Data 2.** Proteomic analysis performed in THP-1 cells transfected with CTRLsi or BAFFsi and treated with IR. **Figure 3—Source Data 3.** Uncropped immunoblots for Figure 3.

Proteomic analysis (**Figure 3—Source Data 2**) revealed several proteins that showed increased abundance in THP-1 cells after IR-induced senescence but were less elevated after BAFF silencing (**Figure 3—figure supplement 1C and Figure 3—Source Data 2**). GO and Reactome pathway analyses confirmed the involvement of BAFF in immune activation and leukocyte degranulation, in promoting ROS production and vesicle trafficking (**Figures 3G, and Figure 3—figure supplement 1C**). The top differentially increased proteins (**Figure 3H,I**) included mediators of inflammation such as NCF1 (Neutrophil Cytosolic Factor 1, also known as p47-phox), Cathepsin B (CATB), and MNDA (Myeloid Nuclear Differentiation Antigen) (Holmdahl et al. 2016; Hannaford, Guo, and Chen 2013; Gu et al. 2022). We also found enrichments in proteins involved in intracellular trafficking like RAB1A, RAB1B, and NSF (Yang et al. 2016; Morgan and Burgoyne 2004); some of these proteins, like NCF1, ACAA2 (THIM), and CYC1 (CY1), are also implicated in oxidation (Holmdahl et al. 2016; Cao et al. 2008; Guo et al. 2017). Analysis of the proteins selectively reduced in senescent cells after silencing BAFF suggested a potential role for BAFF in modulating the organization of the nucleosome and cellular checkpoints, possibly through indirect effectors (**Figure 3—figure supplement 1D-F and Figure 3—Source Data 2**). Integrating the data from both the transcriptomic and proteomic analyses from senescent THP-1 cells uncovered robust roles for BAFF in inflammatory pathways in senescence.

### BAFF promotes SASP in senescent monocytes

The accumulation of senescent cells in tissues with advancing age has been proposed to have detrimental consequences, as they promote age-related declines and diseases (He and Sharpless 2017; Campisi 2013); accordingly, there is much interest in clearing senescent cells for therapeutic benefit (Ge et al. 2021; Nelson et al. 2012). This task is hampered by the intrinsic resistance of senescent cells to apoptosis, and thus we investigated whether BAFF is implicated in the viability of senescent cells by silencing BAFF in THP-1 cells and evaluating cell numbers after treatment with IR. As shown in **Figure 3A**, *BAFF* mRNA was successfully silenced by 72 h after IR (5 Gy), and the levels of cellular BAFF and secreted BAFF (**Figure 4A**, **Figure 4—figure supplement 1A**) followed similar trends. While BAFF silencing slightly reduced the levels of the DNA-damage marker γ-H2AX, we did not observe major changes in cell viability, as determined by analyzing trypan blue exclusion (**Figure 4B**) or by direct cell counting (**Figure 4C**). By 72 h after IR, there was still residual THP-1 cell proliferation that was comparable between CTRLsi and BAFFsi cells (**Figure 4C**). In this regard, BAFF silencing did not affect the levels of p21 (**Figure 4A**), a well-known suppressor of proliferation during senescence and a transcriptional target of p53; in fact, THP-1 cells do not express p53 due to a 26-nucleotide deletion in the p53 coding sequence that prevents p53 production (Sugimoto et al. 1992). Together, these findings indicate that BAFF does not play a major role in the viability or growth of THP-1 cells, and thus it may not be a valuable target for senolysis of senescent monocytes.

**Figure 4.**
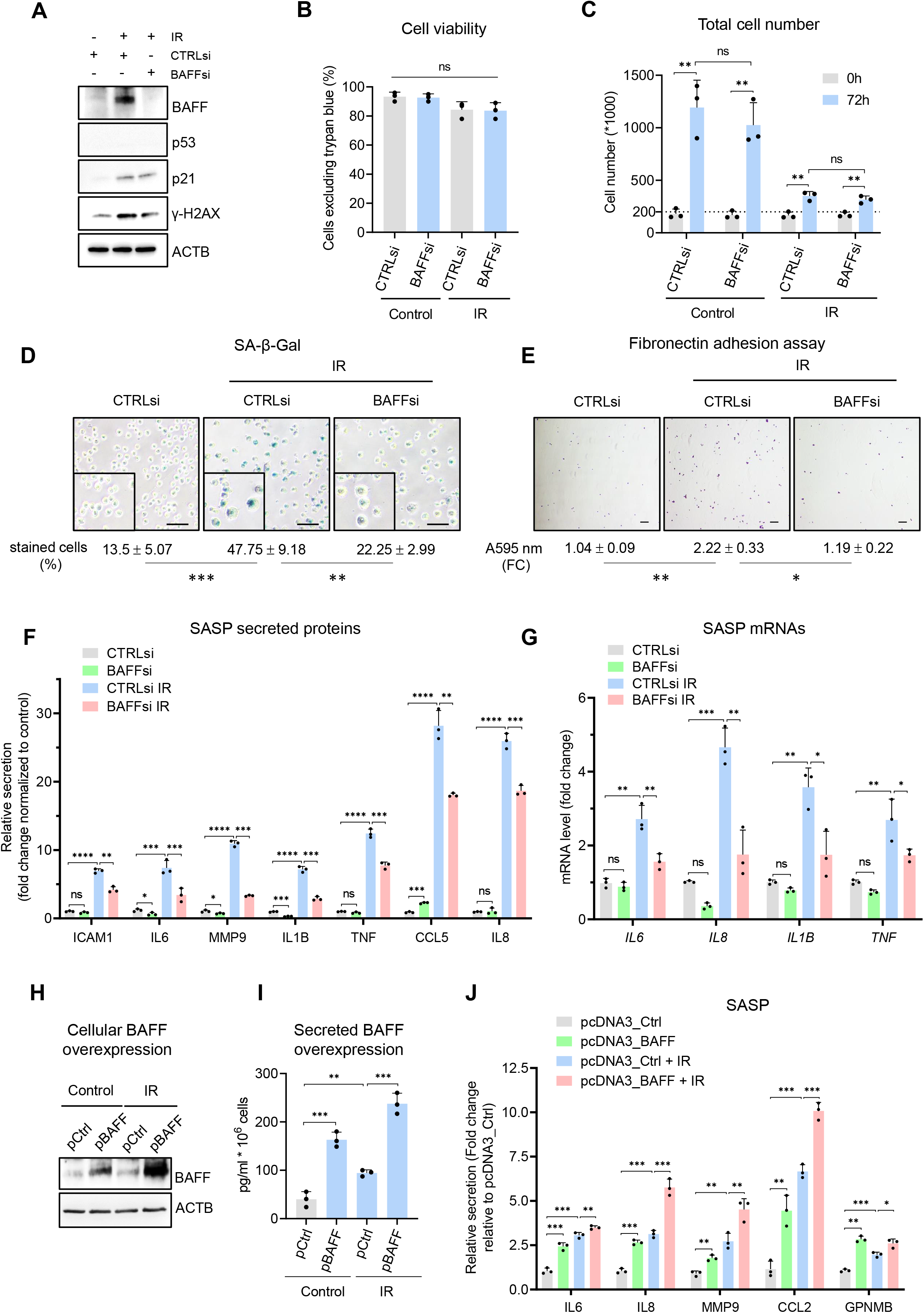
BAFF silencing reduces senescence traits in irradiated monocytes, while BAFF overexpression increases SASP secretion. (**A**) Western blot analysis of the levels of several proteins in THP-1 cells transfected with a control siRNA (CTRLsi) or BAFF siRNA (BAFFsi) following exposure to IR (all the cells were treated with 5 Gy and collected 72 h later); p53 is not expressed in THP-1 cells (Sugimoto et al. 1992); senescence-associated proteins p21 and γ-H2AX were used as positive controls of induced senescence and DNA damage, respectively, and ACTB as loading control. (**B,C**) THP-1 cells were processed as in (A) and cell viability was measured by trypan blue exclusion (% cells excluding trypan blue relative to total cells) (B) and population growth by direct cell counts (C). (**D**) SA-β-Gal activity assay in THP-1 cells processed as in (A); SA-β-Gal-positive cells (% of total cells) were quantified by percentage (%) of positively stained cells as described in Materials and methods. (**E**) Fibronectin adhesion assay showing the presence of adherent activated THP-1 cells (stained with crystal violet) after transfection of CTRLsi or BAFFsi as described in (A). The relative adhesion of monocytes was quantified by crystal violet absorbance (Materials and methods). (**F**) Relative levels (fold change) of secreted cytokines and chemokines in THP-1 cells processed as in (A), as measured by multiplex ELISA 72 h after IR (5 Gy). (**G**) RT-qPCR analysis of the levels of several cytokine and chemokine mRNAs in THP-1 cells treated as in (A) and assayed 72 h after IR. (**H-J**) THP-1 cells were transfected for 16 h with a control plasmid (pCtrl) or a plasmid to express BAFF (pBAFF) and then were either left untreated or treated with IR (5 Gy); 72 h later, whole-cell lysates were studied by western blot analysis (H), and secreted factors by ELISA to detect BAFF (I), and multiplex ELISA assay to detect additional SASP factors (J). Secretion levels are shown as relative fold change compared to the untreated CTRLsi sample. Significance (*p < 0.05, **p < 0.01, ***p < 0.001, ****p <0.0001) was assessed with Student’s *t*-test. Scale bars, 100 μm. Source Data Files for Figure 4: **Figure 4—Source Data 1.** Uncropped western blots for Figure 4.

Given that senescent monocytes display increased adhesion to the extracellular matrix (ECM) and increased secretion of pro-inflammatory molecules (Merino et al. 2011; Elder and Emmerson 2020), we examined if BAFF secreted by THP-1 cells plays an autocrine role in maintaining other traits of THP-1 cell senescence. First, analysis of SA-β-Gal activity revealed that silencing BAFF in THP-1 cells reduced the percentage of SA-β-Gal-positive THP-1 cells (**Figure 4D**). Second, measurement of cell adhesion by using plates coated with fibronectin, a protein component of the ECM that promotes the adhesion of activated monocytes during tissue repair and inflammation (Bauvois et al. 1996; de Fougerolles et al. 2000), indicated lower adhesion of BAFF-depleted as compared to control THP-1 cells (**Figures 4E, Figure 4—figure supplement 1B**). These findings suggested that BAFF might regulate senescence traits associated with monocyte adhesion and possibly the SASP. To test this possibility, we measured a subset of SASP factors in BAFF-silenced THP-1 cells. Interestingly, BAFF silencing led to a marked reduction in the secretion of SASP factors, including typical pro-inflammatory cytokines like IL6, IL8, IL1B, and TNF (**Figure 4F**, **Figure 4—figure supplement 1C**) and their respective mRNAs (**Figure 4G**), supporting the notion that BAFF may promote the SASP in senescent monocytes. To further evaluate if BAFF regulates the SASP, we electroporated THP-1 cells with an empty vector (pCtrl) or BAFF-encoding vector (pBAFF) (Materials and methods) to overexpress BAFF and investigate the changes in SASP. The elevated expression of BAFF, as assessed by western blot analysis and ELISA (**Figure 4H,I**), led to increased secretion of SASP factors, especially IL8, MMP9, CCL2, and CCL5 (**Figure 4J**, **Figure 4—figure supplement 1D**). Furthermore, THP-1 cells displayed small but significant increases in secreted SASP when treated with soluble recombinant human BAFF (60-mer BAFF, Adipogen) (**Figure 4—figure supplement 1E**). These differences were more evident in proliferating cells than in senescent cells, perhaps because ectopic soluble BAFF might only play a limited role in SASP secretion compared to the endogenous or cell membrane-associated BAFF. Overall, BAFF silencing and overexpression experiments suggest that BAFF does not play a role in the viability of senescent monocyte-like THP-1 cells but is important for maintaining the SASP trait.

### BAFF promotes the activation of NF-κB in early senescent THP-1 cells

To study how BAFF promotes inflammation in THP-1 cells, we analyzed the expression of the three known BAFF receptors (BAFFR, TACI, and BCMA) in THP-1 cells. Interestingly, after IR, *BAFFR* and *TACI* mRNAs increased, but *BCMA* mRNA did not (**Figure 5A**). In cells subjected to IR followed by culture for 6 days, western blot analysis indicated that BAFF-R and TACI proteins also increased in senescent cells (**Figure 5B**); additionally, flow cytometry analysis indicated that all three receptors increased on the surface of senescent cells (**Figure 5C**). In sum, the three receptors involved in BAFF signaling via canonical and non-canonical NF-κB pathways (Smulski and Eibel 2018; Afzali et al. 2021; Nagel et al. 2014; Matson et al. 2020), are expressed in senescent THP-1 cells.

**Figure 5.**
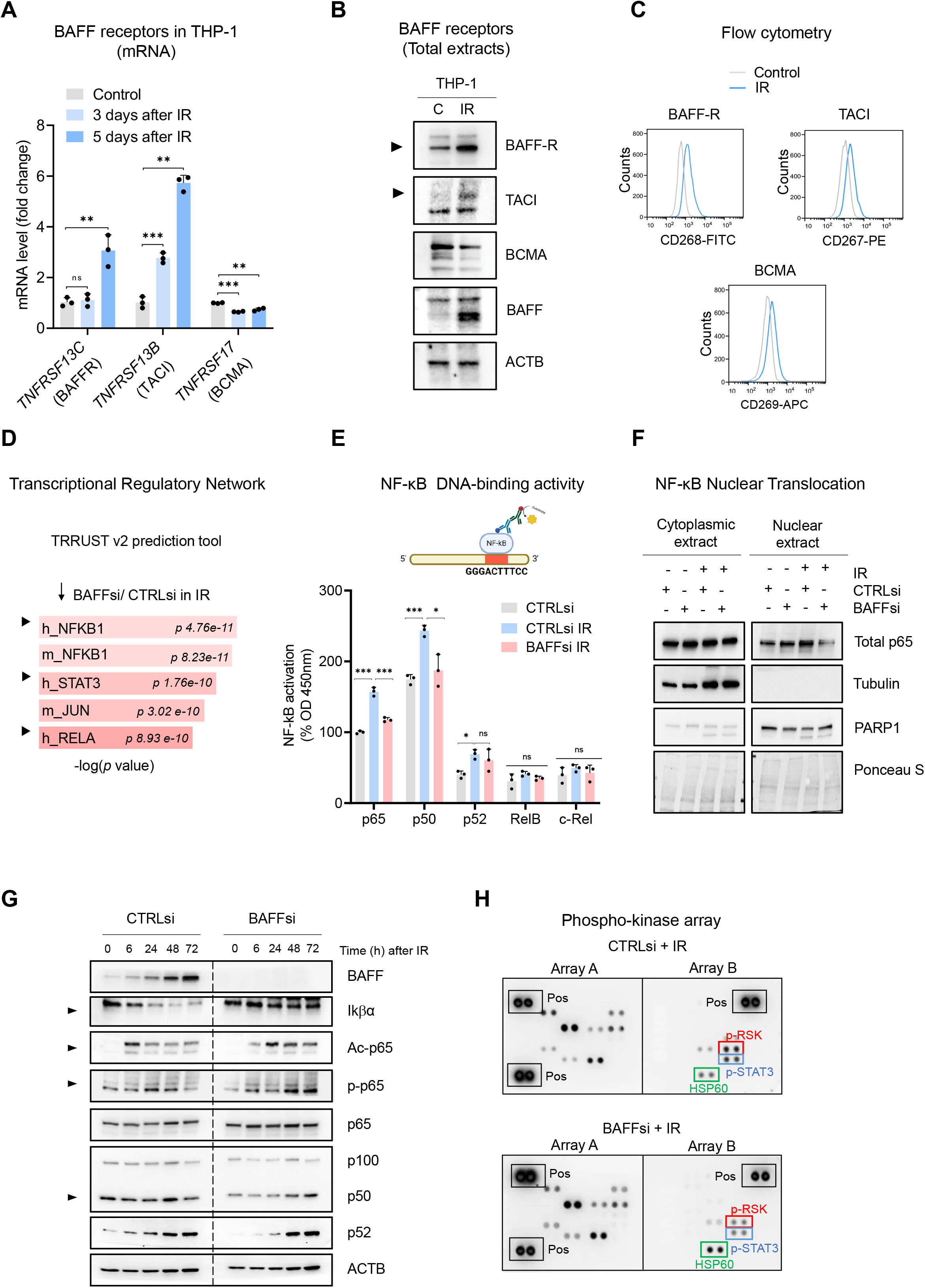
BAFF promotes NF-κB activation in IR-treated THP-1 cells. (**A**) RT-qPCR analysis of the levels of mRNAs encoding the known BAFF receptors in THP-1 cells at 3 and 5 days after treatment with IR (5 Gy). (**B**) Western blot analysis of BAFF receptors in THP-1 cells untreated or irradiated with 5Gy and cultured for 6 days. Due to the low levels of TACI, contrast was increased on the acquired image. (**C**) Flow cytometry analysis of THP-1 cells expressing surface BAFF receptors 6 days after either no treatment or treatment with IR (5 Gy). (**D**) EnrichR analysis of the mRNAs differentially abundant genes obtained from **Figure 3B**. The TRRUST v2 analysis predicts the transcription factors potentially affected by BAFF silencing in THP-1 cells. (**E**) TransAM NF-κB Activation Assay (Materials and methods) performed using THP-1 nuclear extracts to evaluate the binding of different NF-κB subunits to a DNA consensus sequence (*top*), as measured at 450 nm. The basal activity of p65 in the untreated control sample (CTRLsi) was set at 100% and all other values were normalized to it. (**F**) Western blot analysis of p65 levels in nuclear and cytoplasmic fractions of THP-1 cells that were transfected overnight with CTRLsi or BAFFsi and the next day were either left untreated or irradiated and collected 72 h later. Cytoplasmic and nuclear markers (Tubulin and PARP1, respectively) were included to monitor the fractionation procedure; Ponceau S staining served to assess equal loading and transfer. (**G**) Western blot analysis of the proteins in whole-cell extracts prepared from THP-1 cells that were transfected with either CTRLsi or BAFFsi, then exposed to IR (5 Gy) and assessed at the indicated times. Arrowheads point to signals showing differences in NF-κB kinetics after BAFF silencing. (**H**) Phosphoarray analysis of whole-cell lysates (600 μg) prepared from THP-1 cells that had been transfected with CTRLsi or BAFFsi overnight, subjected to IR on the next day, and analyzed 72 h after that. Reference control spots (‘ Pos’) are present on each array. Significance in different panels (*p < 0.05, **p < 0.01, ***p < 0.001, ****p <0.0001) was assessed with Student’s *t*-test. Source Data Files for Figure 5: **Figure 5—Source Data 1.** Uncropped western blots for Figure 5 (part I) **Figure 5—Source Data 2.** Uncropped western blots and arrays for Figure 5 (part II)

EnrichR analysis of differentially expressed transcripts after silencing BAFF suggested a role for BAFF in the regulation of TFs NF-κB and STAT3 in THP-1 cells (**Figure 5D**, **Figure 5—figure supplement 1**). We thus analyzed the DNA-binding activity of multiple NF-κB subunits in senescent THP-1 cells upon BAFF silencing (Materials and methods). IR increased the DNA-binding activity of p65, p50, and p52 in control cells, but not the activities of RelB or c-Rel (**Figure 5E**). Interestingly, the activity of p65 and p50, representing the canonical pathway of NF-κB, was strongly reduced after silencing BAFF (**Figure 5E**). To assess the changes in of p65/RelA activity by other methods, we prepared cytoplasmic and nuclear fractions from proliferating and senescent THP-1 cells expressing either normal levels of BAFF or reduced BAFF levels by silencing. Interestingly, in proliferating THP-1 cells, silencing BAFF did not change the nuclear abundance of p65, while in senescent THP-1 cells, silencing BAFF significantly reduced the levels of nuclear p65 (**Figure 5F**). Tubulin and PARP1 were used as markers for proper cytoplasmic and nuclear fractionation, respectively, and Ponceau S was used to stain the membrane to assess total loaded proteins (**Figure 5F**). These findings suggest a role for BAFF in the canonical pathway of NF-κB activation in senescent THP-1 cells.

A crucial step in the canonical pathway of NF-κB activation is the degradation of the inhibitory IκB proteins, which precedes NF-κB activation and nuclear translocation (Hayden and Ghosh 2008; Israel 2010). Thus, to further study the role of BAFF on the activation of the canonical NF-κB pathway, we analyzed the pattern of IκBα degradation. As shown in **Figure 5G**, IκBα levels declined rapidly after treatment with IR in control cells, but not in cells in which BAFF was silenced (**Figure 5G**), further supporting a role for BAFF in NF-κB activity.

BAFF was previously shown to activate the canonical NF-κB pathway in monocytes after LPS treatment by affecting p65 acetylation, which is required for DNA binding and full transcriptional activity (Gardam and Brink 2014; Lim et al. 2017). Western blot analysis of p65 acetylation relative to total p65 in irradiated THP-1 cells showed a transient peak in p65 acetylation at ~6 h after irradiation in control cells, and a delayed peak at ~24-48 h followed by persistently elevated p65 acetylation in BAFF-silenced cells (**Figure 5G**). Similarly, p65 phosphorylation also showed a delayed peak in BAFFsi cells, reaching a maximum at ~24 h after irradiation in control cells, and at ~48-72h in BAFFsi cells. In sum, the influence of BAFF on IκB degradation, p65 nuclear accumulation and p65 modifications may affect the DNA-binding activity and transcriptional program of NF-κB. Other NF-κB subunits (p50 and p52) changed only modestly as a function of BAFF abundance (**Figure 5G**). Although they may also influence NF-κB activity overall, our results point to a major role of BAFF in the regulation of p65 function.

To assess more broadly other pathways, kinases, and TFs that might be regulated by BAFF in senescent monocytes, we examined the results of phospho-protein array analysis. Interestingly, BAFF silencing in THP-1 cells reduced STAT3 phosphorylation (**Figure 5H**), in agreement with the EnrichR prediction (**Figure 5D and Figure 5—figure supplement 1**) and with the role of STAT3 in SASP production (Kojima et al., 2013). We also observed a reduction in the active forms of RSK 1-3 (**Figure 5H**), previously implicated in the regulation of inflammatory responses and SASP in monocytes undergoing antiretroviral therapy (Singh et al. 2019). Taken together, our findings indicate that BAFF contributes to the activation of proinflammatory pathways in senescent monocytes, at least in part by activating p65/NF-κB and possibly other signaling pathways including STAT3.

### Cell type-dependent roles of BAFF in senescence

To gain a more complete understanding of the role of BAFF in senescence, we investigated its function in primary fibroblasts, which are well-established models for senescence and express the senescence-relevant protein p53. Importantly, WI-38 fibroblasts expressed the mRNAs encoding all three BAFF receptors, BAFFR, TACI, and BCMA (**Figure 6—figure supplement 1A**), and biotinylated protein pulldown followed by western blot analysis confirmed their presence on the surface of WI-38 cells (**Figure 6—figure supplement 1B**); the positive control protein DPP4 (Kim et al. 2017) was also highly expressed on the surface of senescent WI-38 cells. Given that the expression of *BAFF* mRNA and BAFF protein increased more slowly in WI-38 fibroblasts relative to THP-1 cells (see **Figures 2F,2I,5G**, and **Figure 6—figure supplement 1C,D**), we sequentially transfected siRNAs at days 0 (before IR) and 7, and harvested cells at day 10 to achieve a prolonged depletion of BAFF in WI-38 cells (**Figure 6—figure supplement 1E**).

Given that BAFF silencing reduced senescence and the SASP in THP-1 cells (**Figure 4D,F,G**), we tested if fibroblasts displayed similar or different characteristics upon BAFF depletion. As shown, silencing BAFF in senescent WI-38 fibroblasts led to a strong decrease in SA-β-Gal staining (**Figure 6A**), but the effects on the secretion of SASP factors were mainly restricted to changes in IL6 (**Figure 6B** and **Figure 6—figure supplement 1F**). Similarly, BAFF silencing in senescent IMR-90 fibroblasts treated with etoposide (50 μM ETO, followed by 8 days in culture) decreased the SA-β-Gal staining (**Figure 6—figure supplement 1G**) and IL6 expression levels (**Figure 6—figure supplement 1H**), with limited effects on the levels of other cytokines (**Figure 6—figure supplement 1I**).

**Figure 6.**
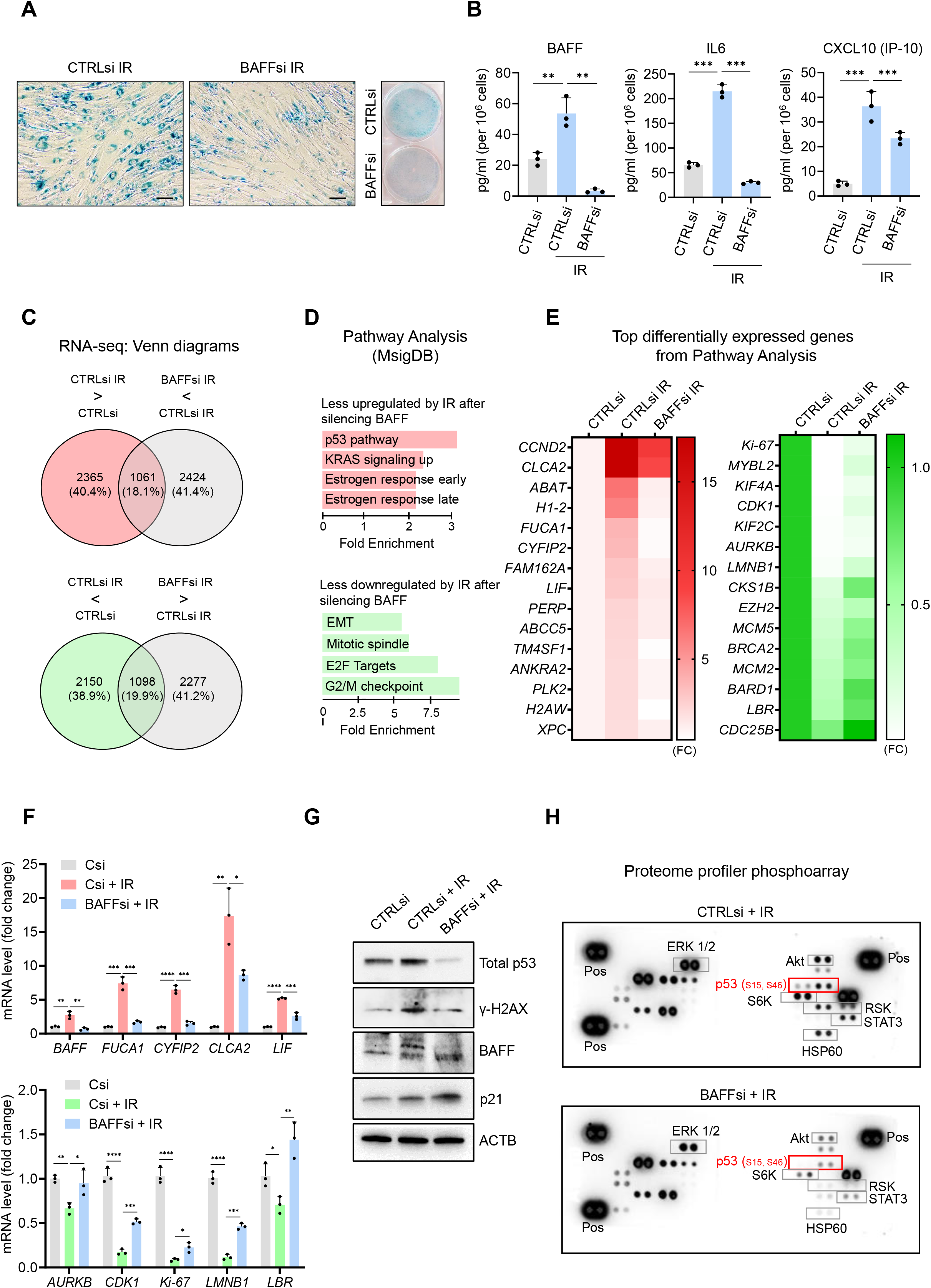
BAFF silencing reduces senescence traits and p53-dependent genes in primary fibroblasts. (**A**) SA-β-Gal activity assay in WI-38 fibroblasts transfected with CTRLsi or BAFFsi, cultured for 18-24 h, treated with a single dose of IR (10 Gy), and assayed 10 days later (Materials and methods, **Figure 6—figure supplement 1E**). Scale bar, 100 μm. (**B**) Levels of BAFF, IL6, and CXCL10 in the culture media of WI-38 cells treated as in (A), measured by multiplex ELISA. Extended data for unchanged cytokines are reported in **Figure 6—figure supplement 1F**. (**C**) WI-38 cells processed as described in (A) were subjected to RNA-seq analysis (GSE213993, reviewer token mbubswukttkhtsf; **Figure 6-Source Data 1**). *Top*, mRNAs showing increased abundance after IR (red) and mRNAs showing reduced abundance after silencing BAFF (gray) are identified at the intersection (Cutoff: padj<0.05, fold change >1.3). *Bottom*, mRNAs showing reduced abundance after IR (green) and higher levels after silencing BAFF (gray) are identified at the intersection (Cutoff: padj<0.05, fold change >1.3). (**D**) MSigDB hallmark analysis performed on the differentially expressed mRNAs in WI-38 fibroblasts. *Top*, pathways less upregulated by IR after silencing BAFFsi. *Bottom*, pathways less reduced by IR after silencing BAFFsi. Bars are ordered according to the fold enrichment of individual pathways. (**E**) Heatmaps of the top differentially expressed mRNAs from the pathway analysis in (D). Data are average of two values and are shown as fold change (FC) relative to the control (CTRLsi FC=1). *Left*, top mRNAs selectively induced by IR and reduced after silencing BAFF. *Right*, top mRNAs selectively reduced by IR but remaining expressed after silencing BAFF. (**F**) Validation by RT-qPCR analysis of representative mRNAs identified by RNA-seq analysis in (C) and listed in the heatmaps in (E). (**G,H**) Whole-cell lysates were prepared from WI-38 cells processed as described in (A); the levels of p53, γ-H2AX, senescence-associated control protein p21, and loading control ACTB were assessed by western blot analysis (G), and the levels of phosphoproteins were detected by phosphoarray analysis (H), which included positive control (‘Pos’) reference dots. Source Data Files for Figure 6: **Figure 6—Source Data 1.**RNA-seq analysis performed in WI-38 fibroblasts transfected with CTRLsi or BAFFsi and treated with IR. **Figure 6—Source Data 2.** Uncropped immunoblots and arrays for figure 6.

To examine in greated detail the role of BAFF in senescent fibroblasts, we analyzed gene expression profiles after silencing BAFF in senescent WI-38 cells. The mRNAs that increased or decreased in senescent WI-38 cells as a function of BAFF levels, as identified by RNA-seq analysis [**Figure 6C and Figure 6—Source Data 1** (GSE213993, reviewer token mbubswukttkhtsf)], highlighted a major role for BAFF in the induction of the p53 pathway (**Figure 6D, *top*, Figure 6E, *left***) and the repression of transcripts involved in cell cycle progression (**Figure 6D, *bottom*, Figure 6E, *right***). RT-qPCR analysis of some of the p53 target genes found in the RNA-seq analysis *(FUCA1, CLCA2, LIF*,and *CYFIP2* mRNAs; **Figure 6F, t*op***) confirmed that their expression increased in senescent cells and was significantly diminished by BAFF silencing. Similarly, RT-qPCR analysis confirmed that the expression levels of many mRNAs (e.g., *AURKB, CDK1, Ki-67, LMB1*, and *LBR* mRNAs) that were significantly reduced in senescent cells were moderately restored by BAFF silencing (**Figure 6E, *right*, and Figure 6F, *bottom***). These data helped to explain the impairment of fibroblast senescence after BAFF silencing. Interestingly, the rise in total p53 levels after IR was strongly reduced upon BAFF silencing (**Figure 6G**), and the reduction in γ-H2AX, a marker of DNA damage, suggested that silencing BAFF might somehow lower genotoxic stress. Analysis of p53 phosphorylation using a phosphoarray further indicated that BAFF silencing also decreased the levels of p53 phosphorylated at Ser15 and Ser46 (**Figure 6H**), highlighting a reduction in transcriptionally active p53. The phosphoarray data revealed wide changes in the phosphorylation of additional proteins after silencing BAFF in WI-38 cells (**Figure 6H**), including reduced phosphorylation of STAT3, which was also seen in THP-1 cells (**Figures 5H, 6H**), that could be explained by the consistent reduction of IL6, an upstream regulator of the STAT3 pathway, across different cell types after BAFF silencing (**Figures 4G, 6B, figure 6—figure supplement 1H,I**). As in THP-1 cells, we also observed reduced phosphorylation of RSK1/2/3 in fibroblasts (**Figures 5H, 6H**).

Overall, our data have uncovered some shared roles for BAFF in senescent monocytes and senescent fibroblasts, including the presence of BAFF receptors, and the loss of both SA-β-Gal activity and IL6 after silencing BAFF. Our transcriptomic analyses (**Figure 6—figure supplement 1J**), as well as western blot analysis of γ-H2AX levels (**Figures 4A, 6G**), suggest a potential role of BAFF in protecting or reducing DNA damage in both THP-1 and WI-38 cells. However, our data also highlight interesting differences in the impact of BAFF on senescence of monocytes and fibroblasts linked to the absence of the senescence regulatory factor p53 in THP-1 cells (**Figures 4A, 5H**) (Fleisher, 2004). By contrast, in senescent WI-38 cells we observed a reduction of p53 after silencing BAFF (**Figures 6G, 6H**), and found that BAFF influenced the phosphoproteome differently in the two cell types (**Figure 5H**, **6H**). Finally, we only observed minor changes in SASP after silencing BAFF in WI-38 cells (**Figure 6B**) while THP-1 cells displayed marked differences in SASP (**Figure 4F**). Together, our results indicate that the senescence-associated protein BAFF is jointly elevated across senescence paradigms but its functional impact on senescence programs varies depending on the cellular context (**Figure 7**).

**Figure 7.**
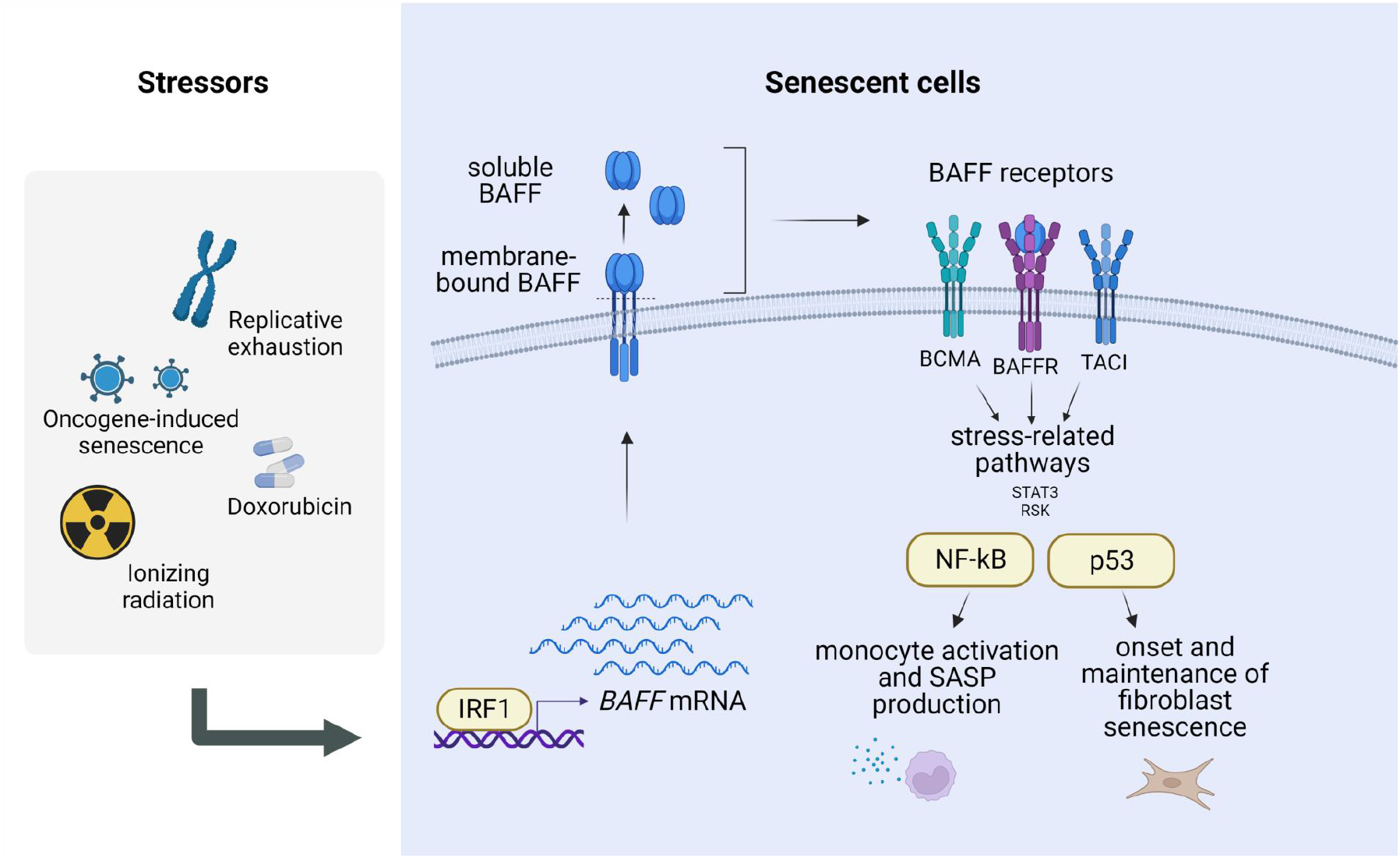
Regulation and role of BAFF in senescence. Model proposed for the regulation and role of BAFF in senescence (created using BioRender). Following DNA damage, the TF interferon-regulated factor IRF1 induces the transcription of *BAFF* mRNA. The protein BAFF is translated and inserted into the plasma membrane, where it can be further processed into a secreted form. Both forms of BAFF are increased in senescence, and both have been previously reported to be functional and capable of activating BAFF receptors (BAFFR, TACI, BCMA), which in turn stimulate stress-related pathways in a cell type-dependent manner, with a predominant activation of the NF-κB pathway in monocytic-like cells, and the p53 pathway in primary fibroblasts. Therefore, BAFF may have pleiotropic actions on senescence-associated phenotypes in different cell types. We propose that BAFF is a novel biomarker of senescence and a regulator of different senescence traits.

## Discussion

Over the lifetime, internal and external factors, like replicative exhaustion, oxidants, viral infection, inflammation, and cancer therapies can cause sublethal cell damage that leads to cellular senescence. Despite their persistent growth arrest, senescent cells remain metabolically active and release a variety of cytokines, chemokines, growth factors, and metalloproteinases, a trait collectively known as the SASP. The negative implications of accumulating senescent cells in tissues is mainly linked to the production of these molecules that promote a pro-inflammatory microenvironment, which in turn activate an immune response, and trigger tissue remodeling with loss of normal tissue architecture. Accordingly, senescent cells are believed to participate in the persistent inflammation that develops with age (‘inflammaging’) and is associated with the development and progression of age-related pathologies like cardiovascular and neurodegenerative diseases, lung and liver dysfunctions, diabetes, and cancer (McHugh and Gil 2018; Munoz-Espin and Serrano 2014; Franceschi et al. 2007).

Despite a heightened interest in cell senescence, the development of translational approaches to identify and clear senescent cells has been hindered by an incomplete understanding of the molecular markers of senescent cells, including those that are universally present in all senescent cells and those that define specific senescent cell subgroups. Here, we focused on BAFF, a cytokine previously predicted to be increased across models of cell senescence in culture (Casella et al. 2019). We found it elevated not only in cultured cell models but also in a model of senescence in mice (**Figure 1**). We propose that BAFF is a novel potential biomarker of senescence both in culture and *in vivo*, as it is easily measurable using sensitive methods like RT-qPCR analysis and ELISA.

BAFF has been extensively studied in immunology, as it plays a primary role in B-cell maturation and survival (Schiemann et al. 2001; Mackay and Browning 2002). Besides its role in the homeostasis of the immune system, BAFF was also implicated in autoimmune diseases like lupus and multiple sclerosis, in part associated with its pro-inflammatory function (Moisini and Davidson 2009; Davidson 2010). We validated the increase of *BAFF* mRNA and protein in multiple senescent models (**Figures 1 and Figure 1—figure supplement 1**) and found that IRF1 transcriptionally elevated *BAFF* mRNA levels in senescence (**Figures 2 and Figure 2—figure supplement 1**). Given that BAFF is mainly produced by monocytes (Yoshimoto et al. 2011; Yoshimoto et al. 2020), we focused on the induction of BAFF in the monocytic cell line THP-1. Transcriptomic and proteomic analyses suggested a role for BAFF in monocyte activation and inflammation in senescence (**Figures 3 and Figure 3— figure supplement 1**), as BAFF depletion led to a striking reduction in the production of SASP factors in irradiated THP-1 cells, while BAFF overexpression or ectopic addition of BAFF increased the secretion of pro-inflammatory molecules (**Figures 4 and Figure 4—figure supplement 1**). THP-1 cells expressed all three BAFF receptors (BCMA, TACI, and BAFF-R), and downstream signaling pathways culminated with the activation of the NF-κB component p65/RelA after DNA damage (**Figures 5 and Figure 5—figure supplement 1**). The activation of this pathway could explain, at least in part, the reduction of SASP factor production after silencing BAFF in THP-1 cells. Interestingly, however, in senescent fibroblasts, BAFF did not have a strong impact on SASP factor biosynthesis, except for IL6 production; instead, BAFF appeared to modulate the levels and activity of the senescence-associated TF p53 (which is not expressed in THP-1 cells). Despite these differences, BAFF did influence key senescence traits in both cell types, including DNA damage, as assessed by monitoring γ-H2AX levels, as well as SA-β-Gal activity and IL6 production (**Figures 4, 6, and Figure 6—figure supplement 1**).

These results agree with the emerging view that senescence is a heterogeneous response across different tissues and cell types, varies according to the inducers of senescence and the time elapsed since senescence was triggered, and is robustly influenced by the microenvironment (Cohn et al. 2022). The discovery of factors shared across senescence paradigms and factors specific for select senescent programs can allow more precise interventions when targeting therapeutically this complex cell population.

To guide future strategies targeting BAFF, it will be important to identify the precise signaling mediators that connect activated BAFF receptors to p53 function in primary senescent cells. It will also be important to study the specific contribution of membrane-bound BAFF relative to secreted BAFF in the implementation of the SASP and other senescence traits. Our preliminary data in THP-1 cells using recombinant BAFF suggest that membrane-bound BAFF has a predominant role over soluble BAFF, as we observed only a minor effect of the recombinant BAFF on the induction of SASP (**Figure 4—figure supplement 1**), although further studies are necessary to confirm this hypothesis. Unexpectedly, however, BAFF-neutralizing agents strongly induced a pro-inflammatory response in THP-1 cells (not shown), supporting a possible reverse signaling triggered by molecules that bind the membrane-bound BAFF (‘ out-to-in’ signaling). However, reverse signaling for BAFF remains a point of debate (Jeon et al. 2010; Nys et al. 2013; Zhang et al. 2015) and further studies are necessary to confirm this hypothesis in senescent cells.

The strategies adopted to eliminate senescent cells or reduce their negative effects include use of senotherapeutic compounds, some of which have entered clinical trials. Senolytic drugs preferentially induce the death of senescent cells over non-senescent cells, while senomorphic compounds modulate the senescent phenotype and SASP production without eliminating senescent cells (Zhang et al. 2015; Niedernhofer and Robbins 2018). Given that BAFF is a SASP factor and plays a key role in modulating senescence-associated phenotypes like the SASP, BAFF could be exploited as a potential target of senomorphic therapy or perhaps even as a marker of the efficacy of senomorphic interventions, not only in laboratory settings in culture, but also in animal models and possibly in human trials. The heterogeneous responses observed between monocytes in fibroblasts (**Figure 7**) underscore the importance of studying the role of BAFF in paradigms of senescence involving other cell types (epithelial cells, myoblasts, adipocytes, endothelial cells, hepatocytes, glial cells, etc.) and different senescence inducers. Specifically, since B lymphocytes have been established as a main target cell for BAFF (Schiemann et al. 2001; Mackay and Browning 2002) it will be critical to study the role of BAFF in senescent B cells and age-related immunosenescence. Future work is warranted to test comprehensively if modulators of BAFF activity have therapeutic value in disease states in which senescent cells are detrimental.

## Materials and methods

### Cell culture and senescence induction

Human acute monocytic leukemia THP-1 (ATCC, TIB-202™) cells were cultured in RPMI-1640 medium (Gibco) supplemented with heat-inactivated 10% fetal bovine serum (FBS, Gibco), and 1% penicillin and streptomycin (Gibco). Human primary WI-38 and IMR-90 diploid fibroblasts (Coriell Institute, AG06814-J, I90-79) were cultured in Dulbecco’s modified Eagle’s medium (DMEM, Gibco) supplemented with 10% FBS, 1% antibiotics, and 1% non-essential amino acids (Gibco). For WI-38 and IMR-90, the karyotype is 46,XX; normal diploid female. human coronary artery vascular smooth muscle cells (hVSMCs) were obtained from LifeLine Cell Technology and were cultured in VascuLife SMC Medium Complete Kit (LifeLine Cell Technology, FC-005). All cultures were maintained in an incubator at 37°C and 5% CO_2_. Senescence was induced by exposure to different doses of ionizing radiation (IR) (5 Gy for THP-1 cells, and 10 Gy for WI-38, IMR-90 and hVSMCs) followed by incubation for the times indicated in text and figure legends. Replicative senescence of WI-38 fibroblasts was achieved by passaging proliferating cells [typically at population doubling level (PDL) of 20-24] until they stopped proliferating (typically PDL >50). Doxorubicin-induced senescence was triggered by treating cells with a single dose of doxorubicin (DOX; 250 nM for WI-38 cells, 10 nM for THP-1 cells) and culturing for the numbers of days specified in each case. Oncogene-induced senescence was induced by transducing the cells for 18 h with a lentiviral vector that expressed RasG12V; lentiviral vectors (control and RasG12V) were a kind gift from Dr. Peter F. Johnson (NCI). The time points at which analyses were carried out are indicated in each experiment. The inhibitors of the interferon pathway Ruxolitinib (Ruxo, used at 1 μM), MRT67307 (MRT67, used at 5 μM), and BX-795 (used at 5 μM) were from InvivoGen.

### Transfection and nucleofection

Silencing interventions in THP-1 and WI-38 cells were performed by transfecting 50 nM of siRNAs using Lipofectamine (RNAiMax, Invitrogen) following the manufacturer’s instructions. Control and BAFF siRNAs were purchased from Horizon Discovery (Non-Targeting siRNA pool Cat. D-001206-14-20 and BAFFsi pool D-017586-01-0005 and D-017586-03-0005). IRF1 pool siRNAs and control siRNAs were from Santa Cruz Biotechnology (sc-35706, sc-37007).

THP-1 cells were transfected at a density of 3×10^5^ cells/ml; 18 h later they were treated with IR (5 Gy) in PBS, given fresh medium, and returned to the incubator. WI-38 cells were transfected at a density of 2×10^5^ cells/well of a 6-well plate; 18 h later, they were treated with 10 Gy IR in PBS, given fresh medium, and returned to the incubator. Plasmid vectors were purchased from GenScript (clone BAFF, OHu22261). Nucleofection (Nucleofector kit V, Lonza, program V-001) following the manufacturer’s instructions was used for overexpression experiments in THP-1 cells. For each reaction, we added 0.5 μg of plasmid to 10^6^ THP-1 cells; 18 h later, cells were treated with 5 Gy IR in PBS, given fresh media, and returned to the incubator. Cells were analyzed at the times specified in the figure legends.

### RNA isolation, RT-qPCR analysis, and RNA-sequencing

RNA was isolated using phenol-chloroform according to the manufacturer’s instructions (TriPure^™^ Isolation Reagent, Sigma-Aldrich). RNA integrity was checked on the Agilent TapeStation using the RNA Screen Tape kit (Agilent). Total RNA (500 ng) was used to calculate mRNA levels by reverse transcription (RT) followed by quantitative (q)PCR analysis. RT was performed by using the Maxima Reverse Transcriptase protocol (Thermo Fisher) and qPCR analysis was carried out using specific primer pairs and SYBR green master mix (Kapa Biosystems) with a QuantStudio 5 Real-Time PCR System (Thermo Fisher). The sequences of primer oligos (from IDT) are listed in **Appendix-Table 1**. Relative RNA levels were calculated by normalizing to *ACTB* or *GAPDH* mRNAs, which encode housekeeping proteins ACTB (β-Actin) or GAPDH, respectively, using the 2^-ΔΔCt^ method.

Sequencing libraries were prepared with TruSeq Stranded mRNA kit (Illumina) following the manufacturer’s instructions. Final libraries were analyzed on the Agilent TapeStation using the D1000 Screen Tape kit and libraries were sequenced on an Illumina Platform with 250 cycles (paired-end, dual indexing). The RNA-seq reads aligned to human genome hg19 Ensembl v82 using Spliced Transcripts Alignment to a Reference (STAR) software v2.4.0j and FeatureCounts (v1.6.4) to create gene counts. Differential gene expression analysis was performed with the DESeq2 package version 1.32.0 (Love et al., 2014) in R (v 4.1.0). The Wald test was used for statistical testing and mRNAs with Benjamini-Hochberg adjusted *p*-values < 0.05 and absolute log2 fold change > 1 were determined as differentially expressed. RNA-seq datasets were deposited in GEO (GSE213993, reviewer token mbubswukttkhtsf).

### SA-β-Gal activity

SA-β-Gal enzymatic activity assay was performed following the manufacturer’s instructions (Cell Signaling Technology). Briefly, adherent cells were washed twice with 1× PBS, fixed for 15 min at 25°C in the dark, and stained in a solution (pH 6.0) freshly prepared following the protocol provided by the manufacturer. Suspension cells (THP-1) were first seeded for 6 h on poly-D-lysine (Gibco)-coated wells before starting the assay. Pictures were acquired by using a digital camera system (Nikon Digital Sight) adapted to a microscope (Nikon Eclipse TS100). SA-β-Gal activity was manually quantified by calculating the percentage of stained cells in three different fields per independent replicate.

### Protein extraction, western blot, proteomics, and surface protein biotinylation and pulldown

To extract total protein, cells were washed twice in cold 1× PBS and harvested in 2% SDS in 50 mM HEPES buffer with freshly added protease and phosphatase inhibitors (Roche). Cell lysates were sonicated and centrifuged for 15 min at 12000 ×*g* to remove the insoluble fraction. Nuclear and cytoplasmic fractions were prepared using the NE-PER kit (Pierce) following the manufacturer’s instructions. The protein content of the cleared lysates was quantified with the BCA assay (Pierce). Lysates were mixed with 4 × SDS Laemmli buffer (Bio-Rad) and boiled at 95°C for 5 min. For electrophoresis through SDS-containing polyacrylamide gels (SDS-PAGE) and western blot analysis, samples were loaded on 4-20% Tris-Glycine gels (Bio-Rad) and transferred onto nitrocellulose membranes using the iBlot kit (Invitrogen). Membranes were blocked for 1 h at 25°C with BSA or milk and incubated overnight with primary antibodies at 4°C. A list of the antibodies used in this study is provided (**Appendix-Table 2**). After washing with 1× TBST, the membranes were incubated with the secondary antibodies for 1 h at 25°C in 5% nonfat milk. After washes, the membranes were incubated with ECL solution (Kwik Quant) before acquiring chemiluminescent signals with a ChemiDoc system (Bio-Rad). Densitometry analysis was performed with ImageJ 1.52A.

To prepare samples for proteomic analysis, 10^7^ THP-1 cells per sample were centrifuged for 5 min at 1000 × *g* and washed twice in 1× PBS. Samples were shipped in dry ice for proteomic MS analysis (Poochon Scientific).

Surface proteins were isolated using the Cell Surface Protein Isolation Kit (Pierce) following the manufacturer’s protocol. Briefly, WI-38 cells were washed with 1× PBS and labeled with a membrane-impermeant Sulfo-NHS-SS-biotin conjugate. Biotinylated proteins were captured on a neutravidin resin, washed and eluted in DTT. The eluted proteins were mixed with sample buffer and prepared for western blot analysis.

### NF-κB activity assay

NF-κB DNA-binding activity was measured using the TransAM NF-κB activity assay (Active Motif). Briefly, 2 μg of nuclear extract containing the activated transcription factors were added to a well coated with a consensus oligonucleotide for NF-κB binding. Samples were incubated for 1 h at 25°C, followed by incubation with primary antibodies that recognized the individual NF-κB subunits (p65, p50, p52, c-Rel, RelB). After incubation with secondary antibodies and signal development, absorbance was read on a microplate reader (Glomax, Promega) at 450 nm.

### ELISA and Luminex assay

The level of BAFF secreted in the cell culture media was assessed with the BAFF hypersensitive soluble human BAFF ELISA kit (sensitivity >8 pg/ml) (Adipogen), following the manufacturer’s instructions, and the plates were read on a Glomax microplate reader (Promega) at 450 nm. The levels of other cytokines and chemokines, as well as the level of cytokines in mouse serum, were assessed by using customized plates (Luminex assay, R&D) and analyzed on a Bio-Plex 200 instrument (Bio-Rad).

### Mice and doxorubicin-induced senescence in vivo

The experimental procedures related to animal work (ASP #474-LGG-2023) were approved by the Animal Care and Use Committee of the National Institute on Aging (NIA/NIH). Mice were imported from The Jackson Laboratory (Bar Harbor, ME) and housed in the animal facility in NIA. C57BL/6 mice at 10 to 12 weeks of age (all females) were treated systemically with doxorubicin to induce cellular senescence in tissues and organs *in vivo*. Briefly, a single dose of doxorubicin (10 mg/kg) and/or vehicle (DMSO) was injected intraperitoneally, and different tissues were collected 14 days later.

### Fibronectin adhesion assay

Fibronectin-coated wells were purchased from Corning. THP-1 cells that had been transfected with CTRLsi or BAFFsi 24 h earlier were irradiated or left unirradiated, and returned to the incubator; 72 h later, 2×10^5^ THP-1 cells were seeded per well on fibronectin-coated wells and incubated at 37°C for 6 h to allow adhesion of activated monocytes. Unbound cells were washed away, while adherent cells were fixed in 4% paraformaldehyde (PFA) and stained in a crystal violet solution (0.1% crystal violet in 5% methanol) for 30 min at 25°C. After extensive washing, the bound and stained cells were solubilized in 10% acetic acid. Absorbance was read with a microplate reader (Glomax, Promega) at 595 nm.

### Immunofluorescence

Adherent cells were washed with 1× PBS and fixed in 4% PFA for 10 min in the dark, washed with 1× PBS and permeabilized with 0.2% Triton-X-100 for 5 min. Fixed and permeabilized cells were washed in 1× PBS and blocked in 10% goat serum (ThermoFisher) for 1 h at 37°C. Primary antibodies were diluted in normal goat serum and incubated with the fixed cells for 18 h at 4°C. After multiple washes with PBS, cells were incubated with secondary antibodies for 1 h at 37°C. After extensive washing, cells were incubated with DAPI (1:5000, ThermoFisher) for 15 min at 25°C in the dark; after the final washes, fluorescent images were acquired with a Keyence microscope. Immunofluorescence on monocytes was performed by first seeding the cells on a layer of Cell-Tak (Corning) for 20 min, following the manufacturer’s instructions. Once attached, cells were processed as adherent cells.

### Flow cytometry

THP-1 cells were pelleted by gentle centrifugation at 500 × *g* for 5 min, followed by washing and resuspension in 1× PBS. Cells were incubated with antibodies directly conjugated to fluorescent dyes (anti-human BAFF-R conjugated to FITC, BioLegend, Cat. #: 316904; anti-human BCMA conjugated to APC, BioLegend, Cat. #: 357505; anti-human TACI, conjugated to PE, Cat. #: 311906, BioLegend) for 15 min at 4°C in the dark. Cells were then washed twice with 1× PBS and flow cytometry analysis was performed on a FACS Canto II flow cytometer (BD Biosciences); data analysis was carried out using FlowJo software (BD Biosciences).

### Cytokine array and phosphoarray

Cytokines and chemokines secreted in the culture media were analyzed with the Proteome Profiler Human Cytokine Array Kit (R&D) following the manufacturer’s protocol. The relative levels of kinase phosphorylation in cell lysates were analyzed with the Proteome Profiler Human Phospho-Kinase Array Kit (R&D) following the manufacturer’s instructions. Chemiluminescent signals from the cytokine array or phosphoarray were acquired on a ChemiDoc machine (Bio-Rad), and densitometry analysis was performed with ImageJ, version 1.52A (NIH).

### Statistical analysis and graphs

Experiments were carried out three times unless otherwise stated. Data were tested for normal distribution and were compared by unpaired Student’s *t*-test, using GraphPad Prism 9. Statistical significance was indicated as follows: *p < 0.05, **p < 0.01, ***p < 0.001, ****p <0.0001. Graphs were generated using GraphPad Prism 9.

## Acknowledgements

We thank Dr. Nikki Noren Hooten and Dr. Michele K. Evans (NIA/NIH) for suggestions throughout this research and for samples. We thank Dr. Peter Johnson (NCI) for providing the lentiviral vectors.

## Funding

This work was funded in its entirety by the National Institute on Aging Intramural Research Program of the National Institutes of Health.

## Legends to Figure supplements

**Figure 1—figure supplement 1. Extended Data from Figure 1.**
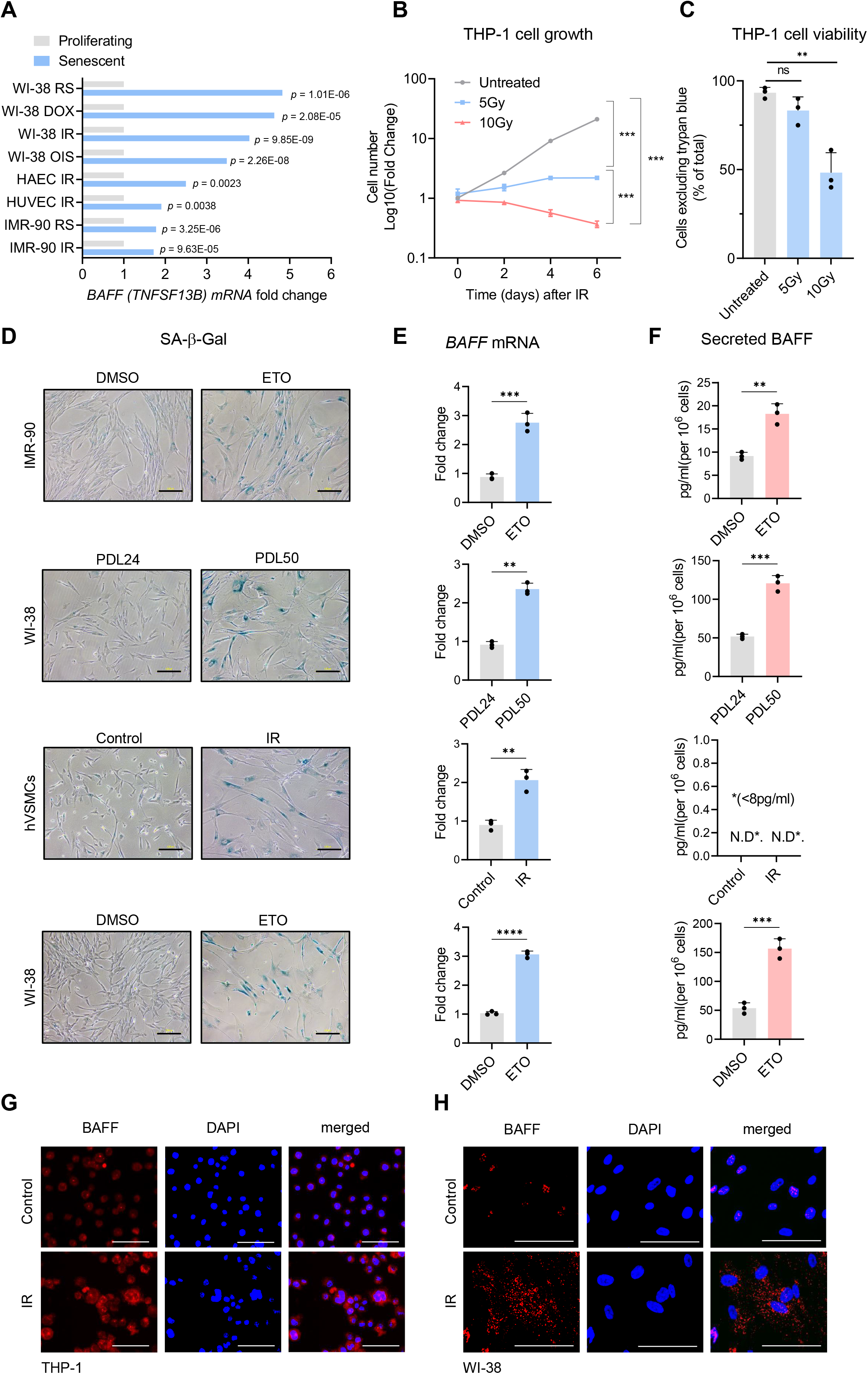
(**A**) *BAFF* mRNA levels across different senescence models; raw data were obtained from our earlier transcriptomic analysis (Casella et al., 2019). (**B**) Growth curve of THP-1 cultures that were left untreated or treated with IR (5 or 10 Gy) and were counted at the indicated times. (**C**) Trypan blue exclusion assay [% cells excluding trypan blue (intact cells) relative to total cells (blue + not-blue)] to evaluate the viability of THP-1 cells treated as in (B) and evaluated at day 6. (**D**) Analysis of SA-β-Gal activity to visualize senescent cells relative to corresponding controls. From top: IMR-90 fibroblasts that were either proliferating (treated with DMSO) or rendered senescent by treatment with a single dose of 50 μM etoposide (ETO) and cultured for 8 days; WI-38 fibroblasts that were proliferating [population doubling level (PDL) 24] or rendered senescent by passaging them until replicative exhaustion (PDL 50); hVSMCs that were either proliferating and otherwise untreated (Control) or were treated with a single dose of IR (10 Gy) and cultured for an additional 7 days; WI-38 fibroblasts that were proliferating treated with DMSO or were treated with a single dose of 50 μM ETO and cultured for 10 days. (**E**) RT-qPCR analysis of *BAFF* (*TNFSF13B*) mRNA levels, normalized to *ACTB* mRNA levels, in the various cells described in (D). (**F**) Levels of soluble BAFF in the culture media of cells treated as in (D) as measured by ELISA. (**G**) Immunofluorescence micrographs to visualize BAFF in THP-1 cells that were rendered senescent by exposure to IR as described in Figure 1A. (**H**) Immunofluorescence micrographs to visualize BAFF in WI-38 cells that were rendered senescent by exposure to IR as described in Figure 1A. Significance (*p < 0.05, **p < 0.01, ***p < 0.001, ****p <0.0001) was assessed by applying Student’s *t*-test. N.D., not detected (<8 pg/ml). Scale bars, 100 μm.

**Figure 2—figure supplement 1. Extended Data from Figure 2.**
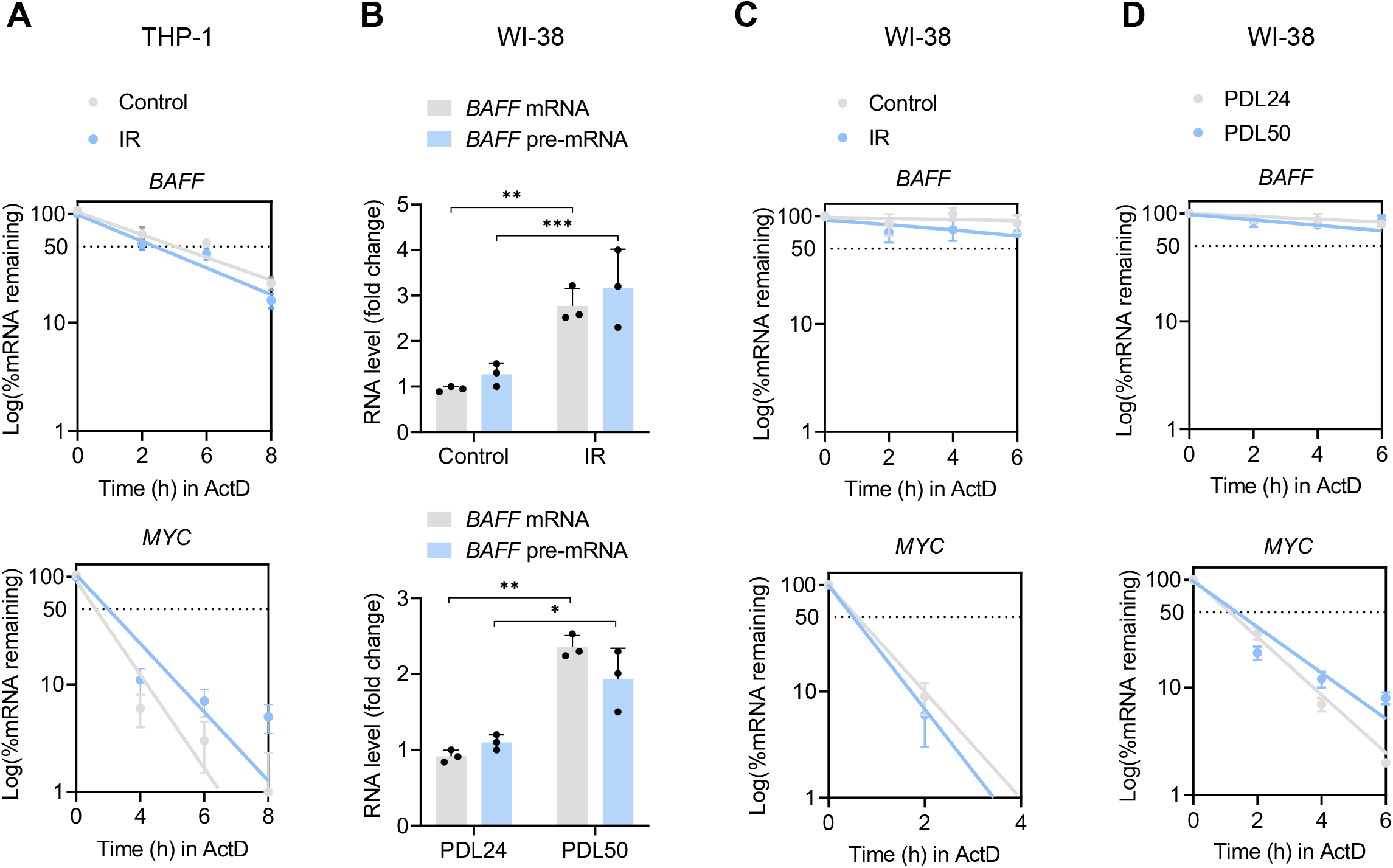
(**A**) Measurement of mRNA stability in THP-1 cells. The levels of *BAFF* mRNA (*top*) and labile control *MYC* mRNA (*bottom*) remaining (% of untreated) after treating cells with the inhibitor of RNA polymerase II actinomycin D (ActD) (2 μg/ml); THP-1 cells were either untreated or had been treated with 5 Gy IR 6 days earlier, and mRNA levels were measured by RT-qPCR analysis of total RNA collected at the times shown after addition of ActD. RNA levels in each sample were normalized to *18S* rRNA levels. Half-lives are estimated as the time needed for each mRNA to reach one-half (50%, discontinuous line) of their abundance at time 0 h. (**B**) RT-qPCR analysis of the levels of *BAFF* mRNA and pre-mRNA in WI-38 cells that were either untreated or treated with a single dose of IR (10 Gy) and cultured for 10 days (*top*) or in WI-38 cells that were proliferating (PDL 24) or replicatively senescent (PDL50) (*bottom*). (**C,D**) Stability of *BAFF* mRNA (*top*) and *MYC* mRNA (*bottom*) as determined following treatment with ActD as explained in (A), in WI-38 cells rendered senescent by exposure to IR (10 Gy) and culture for 10 days (**C**) or by replicative exhaustion (PDL50) (**D**). Significance (*p < 0.05, **p < 0.01, ***p < 0.001) was assessed with Student’s *t* test.

**Figure 3—figure supplement 1. Extended Data from Figure 3.**
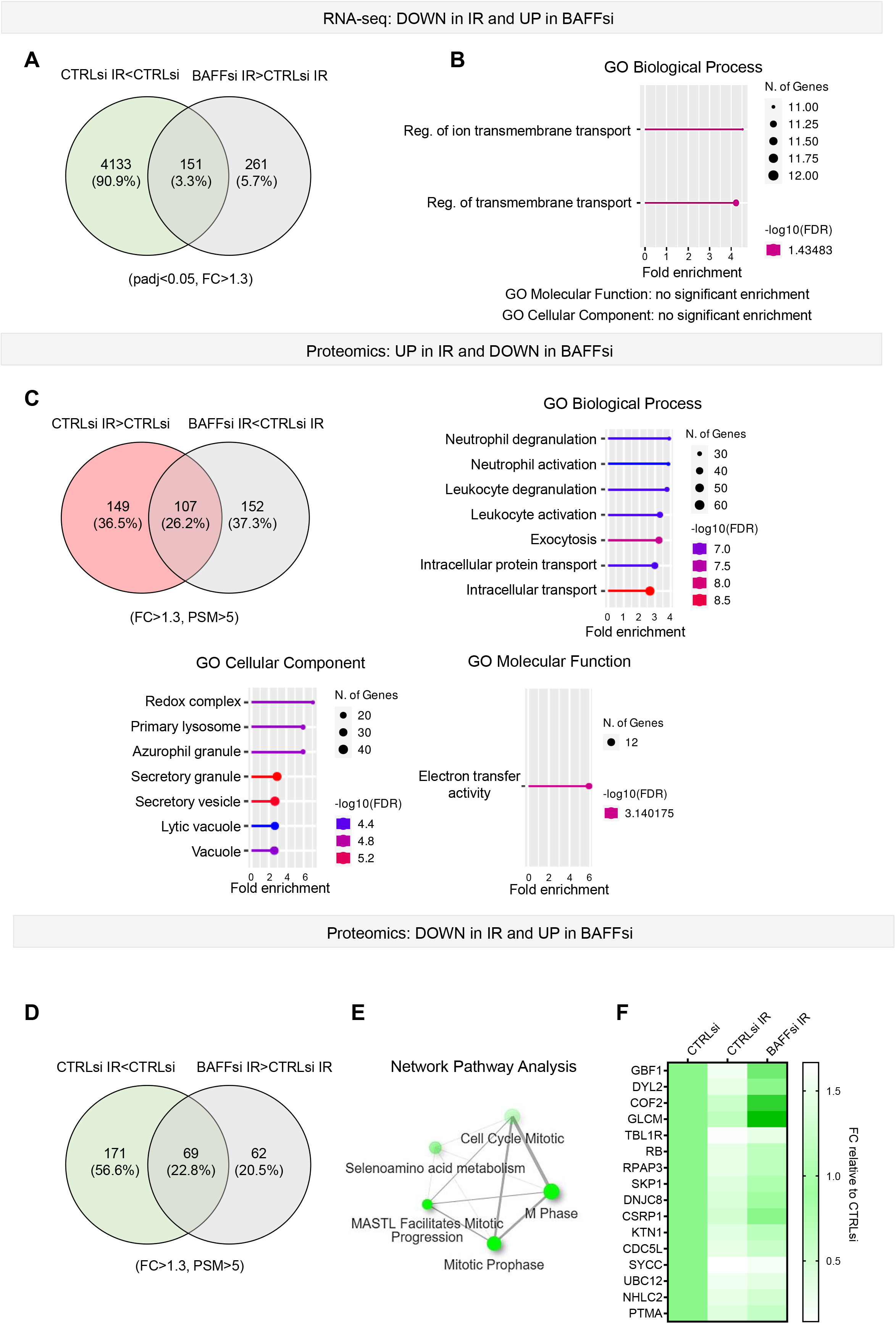
(**A**) Venn diagram showing the mRNAs differentially downregulated in THP-1 cells transfected with control siRNA (CTRLsi) or BAFF siRNA (BAFFsi) upon IR (5 Gy IR, collected 72 h later). Green circle: THP-1 cells exposed to IR (CTRLsi IR) relative to untreated cells (CTRLsi). Gray circle: BAFFsi THP-1 cells exposed to IR relative to CTRLsi THP-1 cells exposed to IR. A complete list of genes from the RNA-seq analysis is available (GSE213993, reviewer token mbubswukttkhtsf, and **Figure 3—Source Data 1**); cutoffs: padj<0.05, [fold change] >1.3. (**B**) Gene ontology analysis (Biological Processes) performed on the differentially downregulated mRNAs identified in (A). GO Molecular Function and GO Cellular Component Diagrams did not show significant enrichments. (**C**) *Top left*, Venn diagram showing the proteins increased after IR (red) and those less upregulated in IR after silencing BAFF (gray) in THP-1 cells. Cells were collected and processed for proteomics as described in Materials and methods. Cutoff: [fold change] >1.3. *Right and bottom*, GO analysis (Biological Processes, Molecular Function and Cellular Component) was performed on the differentially upregulated proteins. (**D**) Venn diagram showing the proteins reduced after IR (green) and those less downregulated in IR after BAFF silencing in THP-1 cells (gray). (**E**) Reactome network analysis highlighting the most highly enriched categories of proteins differentially reduced in CTRLsi or BAFFsi THP-1 cells 72 h after treatment with IR (5 Gy). The plot shows the relationship between the enriched pathways. Two pathways (nodes) are connected if they share 10% or more proteins. Darker nodes are more significantly enriched protein sets; bigger nodes represent larger protein sets; thicker edges represent protein sets with greater overlap. (**F**) Heatmap of the top proteins differentially reduced in BAFFsi IR and CTRLsi IR. Data are shown as fold change between PSM relative to the control (CTRLsi: FC=1). Cutoff: [FC]>1.3, proteins with detected PSM score above 5. Diagrams in (B, C, E) were created with ShinyGO.

**Figure 4—figure supplement 1. Extended Data from Figure 4.**
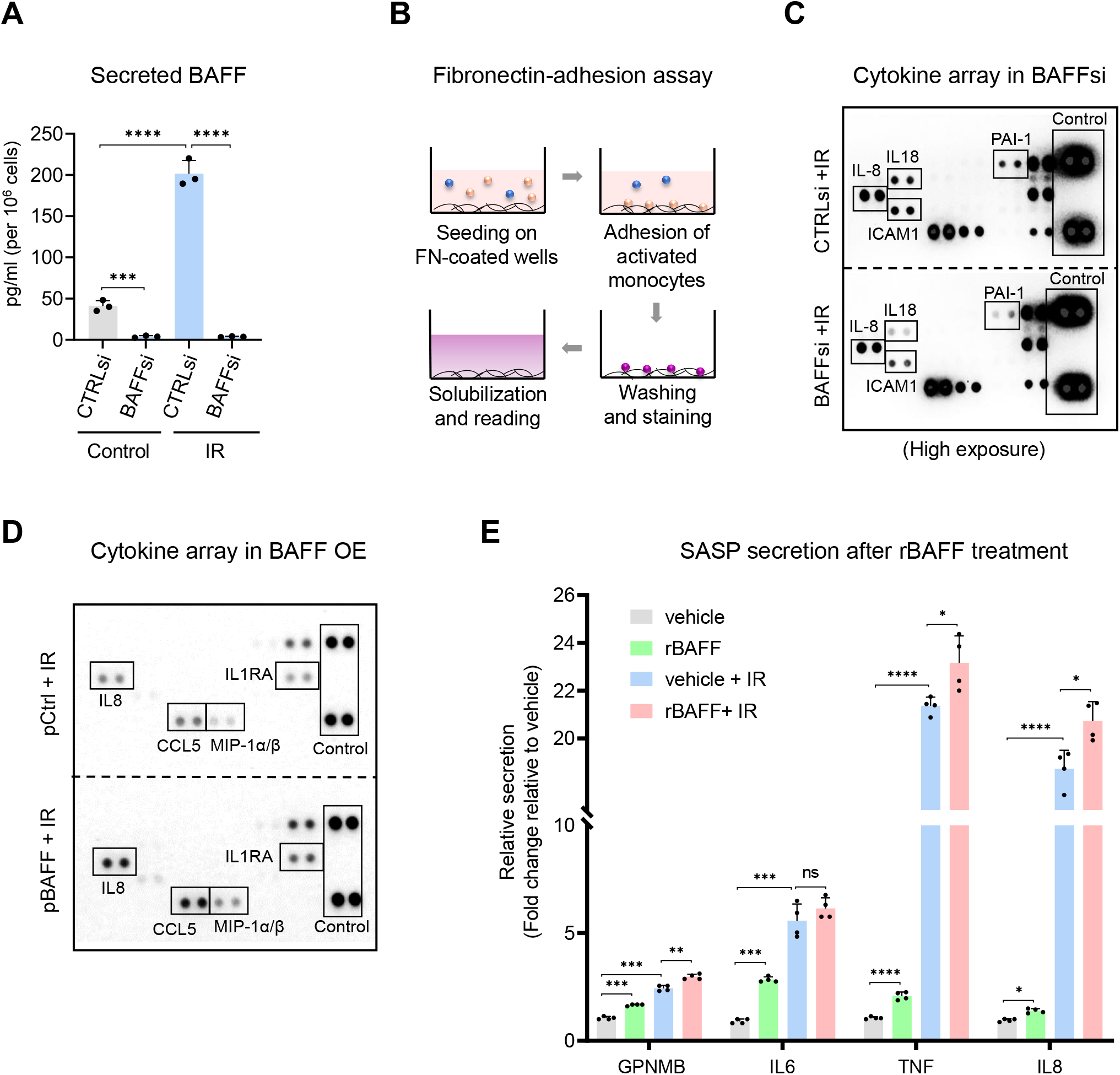
(**A**) THP-1 cells were transfected with CTRLsi or BAFFsi; 18 h later, they were irradiated with 5 Gy and 72 h after that they were analyzed by ELISA. (**B**) Schematic of the fibronectin adhesion assay (Materials and methods) used in **Figure 4E**. (**C**) Cytokine array performed using the media of THP-1 cells treated as in (A). The high exposure was chosen to appreciate differences in low-abundance proteins like IL18 and PAI-1. Positive control spots are present on each individual array. Media were collected from 2×10^6^ cells per sample. (**D**) Cytokine array performed on the media of THP −1 cells transfected overnight with Control (pCtrl) or BAFF (pBAFF) plasmid, treated with IR (5 Gy) on the next morning and assayed 72 h later. Reference control spots are present on each individual array. Media were collected from 2×10^6^ cells per sample. (**E**) Multiplex ELISA to measure secreted factors in THP-1 cells that were treated with either vehicle (DMSO) or 200 ng/ml human recombinant 60-mer BAFF (Adipogen), in untreated cells or irradiated cells (5 Gy) immediately after irradiation and up to 72 h. Significance (*p < 0.05, **p < 0.01, ***p < 0.001, ****p <0.0001) was assessed with Student’s *t-*test. Source Data for **Figure 4—figure supplement 1**: **Figure 4—figure supplement S1—Source Data 1.** Uncropped blots and array for **Figure 4— figure supplement 1**.

**Figure 5—figure supplement 1. Extended Data from Figure 5.**
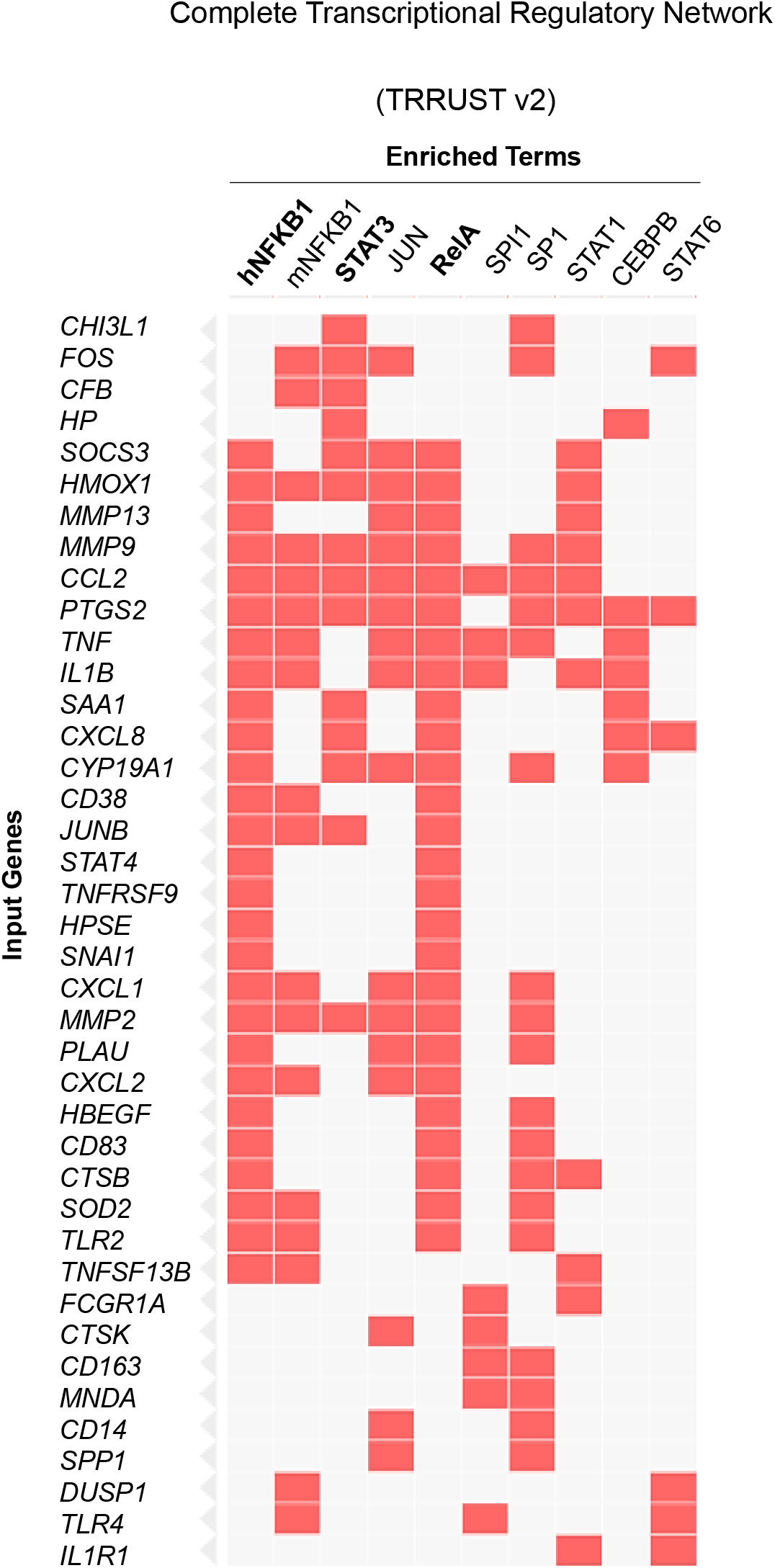
Complete EnrichR analysis performed on the differentially expressed genes from **Figure 3B**. The analysis predicts the transcription factors potentially affected by BAFF silencing in THP-1 cells. Each column shows the individual genes under the control of a specific transcription factor.

**Figure 6—figure supplement 1. Extended Data from Figure 6.**
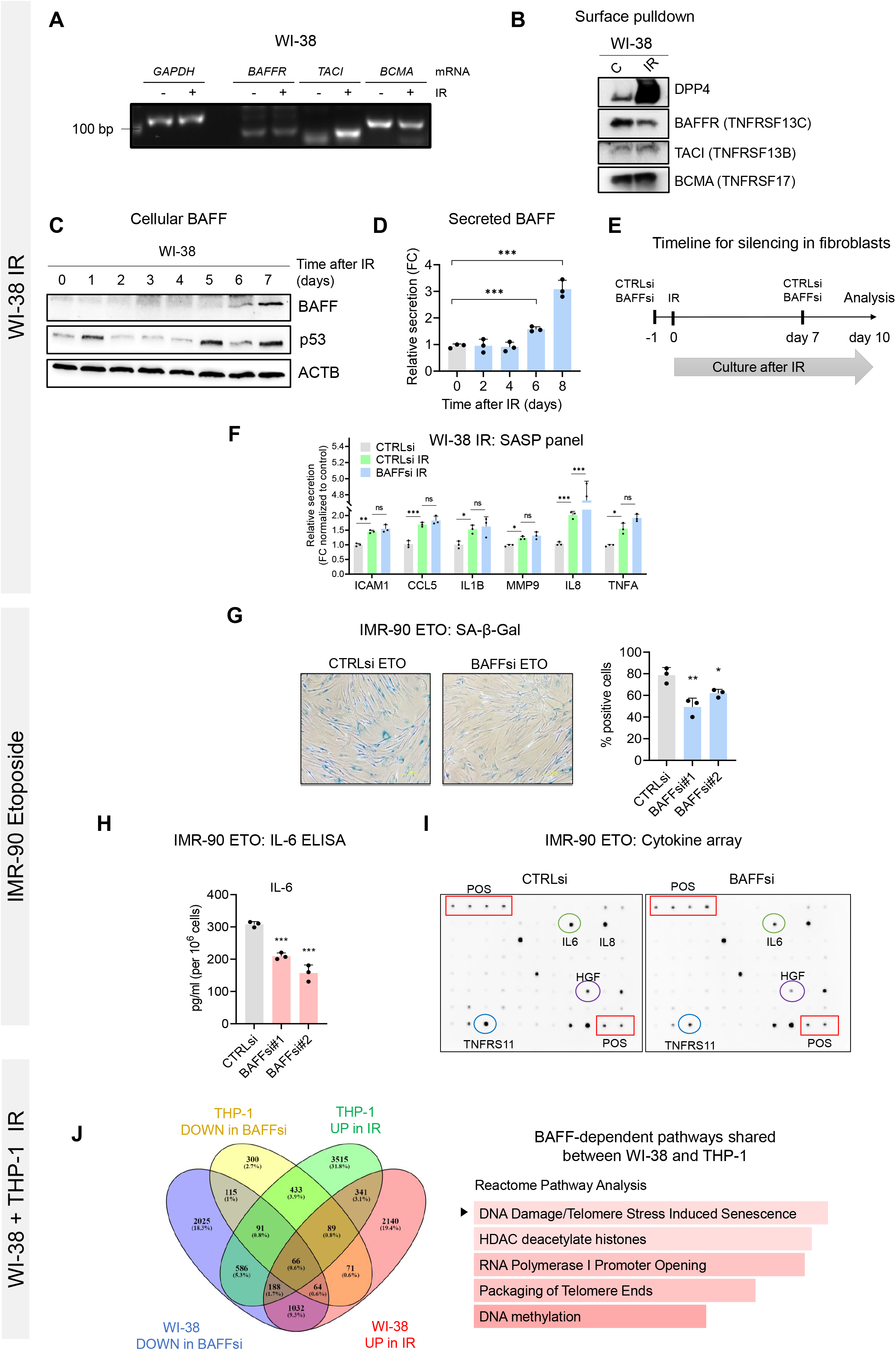
(**A**) RT followed by conventional PCR analysis to visualize the mRNAs encoding the three BAFF receptors in WI-38 cells that were treated with IR (10 Gy) and cultured for an additional 10 days before RNA extraction and analysis. PCR amplicons were visualized after electrophoresis on a 2% agarose gel; *GAPDH* mRNA amplicons were assayed to monitor differences in sample input. (**B**) Pulldown of biotinylated surface proteins and western blot analysis showing the expression of the BAFF receptors in WI-38 cells treated as in (A); detection of DPP4 served as a positive control for senescence and a marker of surface proteins. (**C,D**) WI-38 cells were treated with IR (10 Gy) and the levels of BAFF, total p53, and loading control ACTB were assessed by western blot analysis (C) and the levels of soluble secreted BAFF were assessed by ELISA (D) at the times indicated. (**E**) Schematic of the timeline of BAFF silencing and exposure to IR in WI-38 cells. (**F**) Extended data relative to the multiplex ELISA in **Figure 6B**. **(G)** IMR-90 fibroblasts were transfected with CTRLsi or BAFFsi, and 24 h later they were treated with 50 μM etoposide (ETO) and cultured for a total of 8 additional days, whereupon SA-β-Gal activity was assayed. Scale bar, 100 μm. Graph, quantification of SA-β-Gal (% positive cells). **(H,I)** IMR-90 cells were treated as in (G) and the culture media were used to quantify the levels of IL6 by ELISA (H) and the levels of multiple secreted factors by cytokine array analysis (I). (**J**) *Left:* Venn diagram showing the number of BAFF-responsive genes shared between CTRLsi and BAFFsi THP-1 and WI-38 cells. Data were obtained by combining the upregulated genes from the RNA-seq analysis (GSE213993, reviewer token mbubswukttkhtsf) and **Figure 3—Source Data 1, Figure 6—Source Data 1**. Cutoff: padj<0.05; [fold change] > 1.3. Diagram was created with Venny 2.1.0. *Right:* Reactome pathway analysis of the data in the Venn diagram, and represented using ShinyGO; bars are ordered according to the fold enrichment. Significance (*p < 0.05, **p < 0.01, ***p < 0.001, ****p <0.0001) was assessed with Student’s *t* test. Source Data for **Figure 6—figure supplement 1**: **Figure 6—figure supplement 1—Source Data 1.** Uncropped blots and array for **Figure 6—figure supplement 1.**

## Notes

### Competing Interest Statement

The authors have declared no competing interest.

